# Discovery and characterization of a first-in-field transcription factor BRN2 inhibitor for the treatment of neuroendocrine prostate cancer

**DOI:** 10.1101/2022.05.04.490172

**Authors:** Daksh Thaper, Ravi Munuganti, Adeleke Aguda, Soojin Kim, Shengyu Ku, Olena Sivak, Sahil Kumar, Sepideh Vahid, Dwaipayan Ganguli, Himisha Beltran, Colm Morrissey, Eva Corey, Amina Zoubeidi

## Abstract

The increased incidence of treatment-emergent neuroendocrine prostate cancer (NEPC) is particularly alarming as this diagnosis is associated with poor prognosis. Despite initial responses to platinum-based chemotherapy, relapses are common and there is no effective second line therapy for NEPC. We previously identified that neuronal transcription factor BRN2 (POU3F2) is a potent driver of neuroendocrine differentiation and an attractive target for NEPC. Utilizing a combination of *in silico* modeling and X-ray crystallography followed by structure-based lead optimization, we have developed the first potent, specific and orally bioavailable BRN2 inhibitor (B18-94), which inhibits the interaction between BRN2 and DNA. This loss of BRN2 on the chromatin drastically reduces its transcriptional output resulting in downregulation of several known targets in NEPC such as *SOX2, ASCL1* and *PEG10*. Additionally, B18-94 reduces specifically cell proliferation specifically in multiple NEPC models with no effect on adenocarcinoma and other BRN2 negative prostate cancer models. Importantly, the consistency in the transcriptomic changes driven by B18-94 and or CRISPR/Cas9 mediated BRN2 knockout confirmed the on-target specificity, with both methods of BRN2 inhibition downregulating pathways involved in cellular plasticity and proliferation. Finally, we have demonstrated that B18-94, the first-in-field POU-domain transcription factor inhibitor, significantly reduced tumor growth in several NEPC xenograft models with no observable toxicity, suggesting potential for therapeutic intervention of NEPC.

## Introduction

Neuroendocrine prostate cancer (NEPC) is an aggressive histological variant of prostate cancer (PCa). This variant represents approximately 1% of new prostate cancer diagnoses^1^. However, with implementation of newer more potent androgen receptor signaling inhibitors (ARSI) such as abiraterone and enzalutamide (ENZ) the occurrence of treatment-emergent NEPC (tNEPC) is close to 20%.^2^ Cytotoxic chemotherapies remain the standard-of-care for patients with NEPC and despite initial responses, relapses are common and prognosis is measured in months. Therefore, effective therapies for this deadly disease is an urgent clinical need. Using a unique model of enzalutamide-resistance (ENZR), we previously identified that the neuronal transcription factor (TF) BRN2 (*POU3F2*) as a crucial regulator for the development and maintenance of tNEPC^3^ is highly expressed in both de novo NEPC and tNEPC tumors^2,3^. Not limited to NEPC, BRN2 is significantly upregulated in small cell lung (SCLC) and bladder cancers.^4,5^ Targeting BRN2 in SCLC models reduced cell proliferation and neuroendocrine marker expression.^6^ As a key developmental TF for neuronal cell lineage in the hypothalamus^7^, BRN2 has been extensively studied for its ability to induce neurons in both mouse and human lineages in co-operation with other TF like *ASCL1* and *SOX2*.^8,9^ Furthermore, BRN2’s expression is paramount for maintaining the tumor-initiating potential of glioblastoma cells^10^ and it is a critical driver of invasive and metastatic melanoma^11^.

Genetic targeting of BRN2 reduces cell growth both in vitro and in vivo in many tumor types and presents an especially attractive target, especially for small cell neuroendocrine tumors. Historically, targeting of TFs (like AR) has typically been focused on the ligand binding domains (LBD); and small molecules inhibiting TFs that lack LBDs are exceedingly rare. As a member of the POU-domain family transcription factors, BRN2 contains the POU_S_ and POU_H_ DNA binding domains (DBD) held together by a small but flexible linker sequence,^12^ which provides an attractive pocket for a small molecule. Herein, we describe the development of scaffold B18 its derivative, B18-94, a small molecule with enhanced potency and promising pharmacokinetic properties creating a potent chemical biology tool to study BRN2 inhibition in NEPC.

### Discovery and Validation of BRN2 inhibitor

POU-family TFs have a highly disordered N-terminus and no LBD.^12^ In the absence of human BRN2 crystal structure, we used homology modeling to build a high-quality structure of the BRN2-DBD **(Fig. S1A)**. Using SiteFinder within MOE,^13^ we identified a solvent exposed groove between POU_H_ and POU_S_ domains of BRN2 as a potential binding site for small molecules **(Fig. S1B)**. The polar residues around the periphery of the pocket (C299, Q305 and T306) allows for electrostatic interactions while the valine rich core (V396, V397 and V400) provides additional hydrophobic links. Taking the advantage of this structural information, a virtual library of four million small molecules (ZINC v15) was docked into BRN2-DBD. Top 2000 compounds with highest predicted binding affinities were evaluated to exclude reactive/toxic motifs. Adjusting for chemical diversity, seventy-two compounds **(Table S1, Fig. S1C similarity tree)** were subjected to series of in vitro selection steps via reporter assay (luciferase activity), binding to full-length BRN2 by Drug Affinity Responsive Target Stability (DARTS) assay and cell-killing of BRN2 positive cells **(Fig. S1C)**. The selection pipeline identified two compounds: B7 (dihydroindene core) and B18 (benzodiazole core). In this study, we report the lead optimization and activity profile of the B18.

In parallel to medicinal chemistry optimization, we successfully solved the X-ray crystallographic structure of BRN2-DBD (residues 264 – 415) in complex with the More of PORE (MORE) DNA sequence. The refined model with no Ramachandran outliers and an average B factor of 24.2 resulted in a structure solved with 1.9Å resolution **(Fig. 1A, Table S2)**. This structure was highly comparable to the *in silico* model with RMSD of 3.1 **(Fig. S1D)**. As expected, DNA was sandwiched between the two POU DBDs with helix 3 of both domains docking into the major groove of DNA and the N-terminal arm of the POU_H_ domain docking into the minor groove. Interactions with DNA were detected at Q305, T306, R310, S317 of the POU_S_ domain and T359, I361, R399, N404 of the POU_H_ domain.

**Figure 1:**
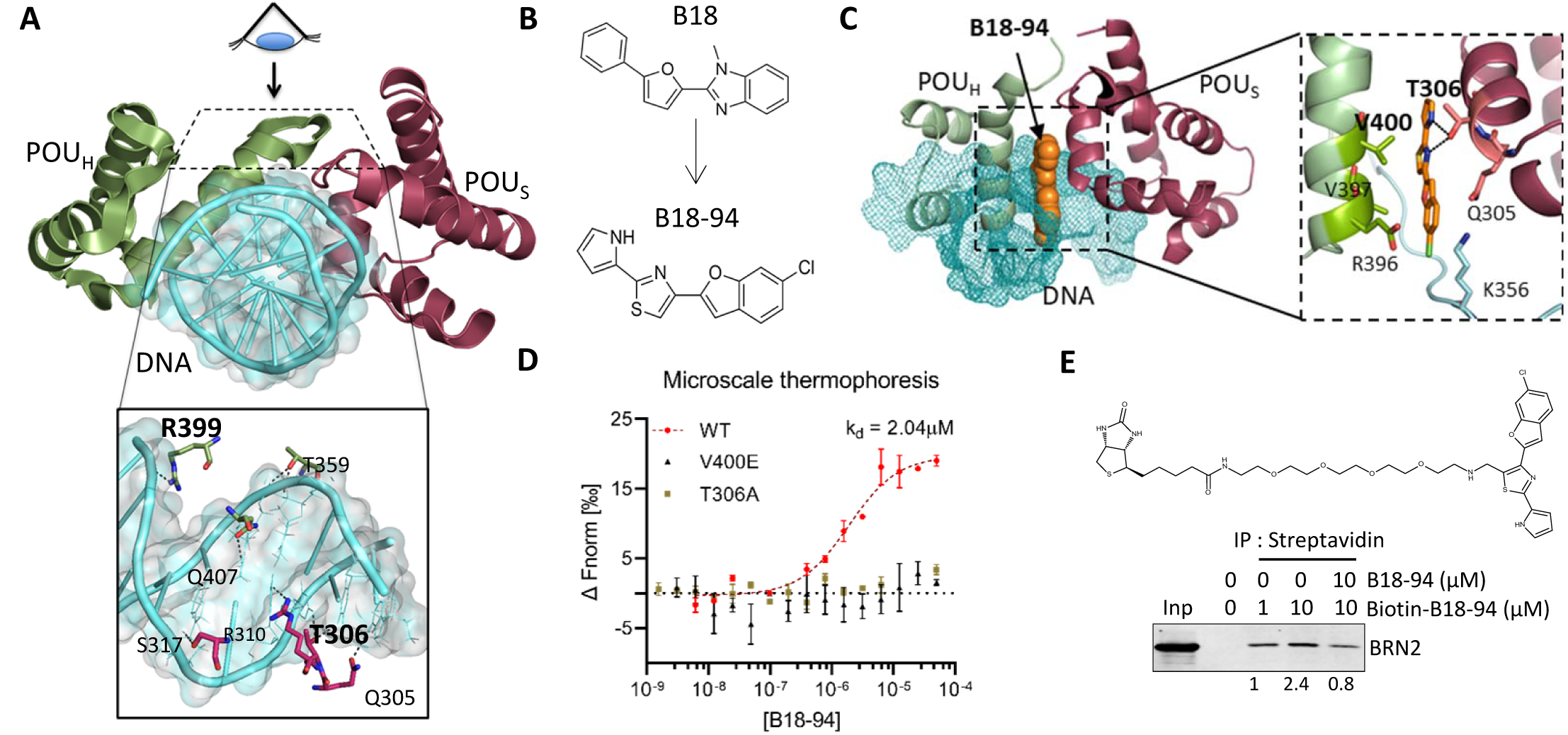
Discovery and validation of BRN2 inhibitor. **(A)** Crystal structure model of BRN2 DNA-binding domain with MORE DNA sequence at 1.9Å. **(B)** Med-chem evolution of compound B18 to B18-94. **(C)** Resultant structure from *in silico* molecular dynamics simulation of 100ns of BRN2 DNA-binding domain with B18-94 with overlay of MORE DNA from crystal structure. **(D)** Microscale Thermophoresis for B18-94 against purified BRN2^WT^, BRN2T^306A^ and BRN2^V400E^. **(E)** Structure of B18-94 with linker and biotin upper panel. Immunoprecipitation with Streptavidin with increasing doses of Biotinylated-B18-94 (1µM and 10µM) competed with 10µM of B18-94 followed by western blot using BRN2 antibody.

Optimization of scaffold B18 through structure-activity relationship established by *in vitro* characterization yielded a more stable and potent derivative, B18-94 **(Fig. 1B, S2A-C)**. To understand the interactions between BRN2 and B18-94, we conducted exhaustive crystallography studies. Across hundreds of solubility matrices, we discovered that due to incompatibilities, the protein-inhibitor complex is not amenable to crystallographic analysis. Instead, we conducted explicit solvent molecular dynamics (MD) simulations for 100ns using AMBER^14^ and through cluster analysis on the MD trajectories identified the most probable binding pose of B18-94 in the BRN2-DBD region **(Fig. 1C)**. The relatively stable RMSD for the final 50ns of the MD simulation indicates that B18-94 maintained its binding orientation, thus achieving its most probable confirmation within the BRN2-DBD **(Fig. S3A)**. This *in silico* conformation predicts that that B18-94 interacts with T306, K356 and V400 of the BRN2-DBD **(Fig. 1C, S3B)** overlapping (T306) or adjacent (V399) to the exact amino acid residues that interact with DNA **(Fig. 1A)**. Supporting these predictive models, microscale thermophoresis (MST) revealed that BRN2^WT^ interacts with B18-94 with a Kd of 2.04 µM **(Fig. 1D)** and the binding was further validated by in-cell DARTS assay **(Fig. S3C)**. We also tested the interaction between B18-84 and BRN2 mutants with MST and DARTS assay. Our data revealed that B18-94 did not bind to BRN2^T306A^ nor BRN2^V400E^ mutants, further supporting the hypothesis that the binding of B18-94 is mediated by these residues **(Fig. 1D, S3D)**. Moreover, biotinylated B18-94 pulled down BRN2 from NEPC cells in a dose dependent manner, which could be antagonized using naked B18-94 **(Fig. 1E)**. These data demonstrate that B18-94 interacts with the BRN2-DBD.

### B18-94 inhibits BRN2 activity in NEPC models

BRN2 protein **(Fig. S4A)**, B18-94 IC_50_ and transcript levels all the POU family members across a panel of PCa cell lines were determined to test the specificity of B18-94. Mapping the IC_50_ of B18-94 to the transcript levels demonstrates a strong negative and statistically significant correlation between BRN2 expression and IC_50_, a trend that was specific to BRN2 **(Fig. S4B-C)**. In addition to NEPC cell lines 42D^ENZR^ (previous characterized by our lab^3^) and NCI-H660, 22RV1 (a common CRPC model), displayed a high level of BRN2 expression and low IC_50_ for B18-94. Interestingly, 22RV1 cells express very low levels of prostate specific antigen (PSA or *KLK3*) with loss of canonical AR signaling and enrichment of neuronal pathways as well as many NEPC markers **(Fig. S5A-B)**; an observation corroborated by Labrecque et al. 2021^15^. The low AR activity, combined with high expression of BRN2 and NEPC splicing factor SRRM4 leading to functional loss of RE-1 silencing TF (REST)^16^ **(Fig. S5C)** are all factors that contribute to the high levels of neuronal signaling in 22RV1 cells. Overall, B18-94 significantly reduced cell proliferation over-time in all three models with high BRN2 expression compared to PCa cell lines lacking BRN2 expression **(Fig. S5D-E**) and increased apoptosis in the BRN2 expressing cell models **(Fig. S5F)**.

In order to evaluate the on-target effect of the BRN2 inhibitor, we compared the effect to BRN2 knockout (BRN2^-/-^) using an inducible CRISPR/Cas9 system **(Fig. S6A)** to effects of B18-94 in both 42D and 22RV1 cells. With the knockout confirmed by western blot, quantitative RT-PCR results showed that change in NEPC marker expression was comparable in 42D^ENZR^ and 22RV1 cells treated with B18-94 to that of the BRN2^-/-^ cells **(Fig. S6B-C)**. Overall, treatment with B18-94 downregulated both protein and mRNA levels of BRN2 itself and several of the known NEPC drivers/targets, e.g., ASCL1,^17^ SOX2,^18^ EZH2,^19^ PEG10^20^ and the NEPC marker CHGA in all 3 models **(Fig. S6A-E)**. We next performed RNA-seq using 42D^ENZR^ cells treated with B18-94 or with BRN2^-/-^ **(Table S3)** and plotted the fold-change (FC) versus control for both conditions. We observed a significant correlation between the directionality of relative gene expression changes (up- or down-regulation) upon inhibition of BRN2 by B18-94 or BRN2^-/-^ **(Fig. 2A)**. Out of 1984 genes with -1 > log2FC > 1, the expression 1654 (84%), genes were concordant between both conditions. Upregulated genes (Q1) were enriched for epithelia development and apoptosis pathways, while downregulated genes (Q3) were enriched for cell cycle and neurogenesis pathways **(Fig. 2B)**. Taken together, these data reveal a consistent reduction in cell proliferation as well as a switch from a neuronal to an epithelial phenotype upon either method of BRN2 inhibition. This high concordance led to a large overlap in biological clusters across gene sets. For example, there was a consistent downregulation of cell cycle/proliferation pathways, cell plasticity pathways related to neuronal differentiation, cancer stem cell (CSC) phenotype as well as the up-regulation of apoptosis and stress response pathways **(Fig. 2C, S7A)**. Notably, RNA-seq further demonstrated the comparable downregulation of NEPC drivers/targets (e.g. ASCL1, SOX2, EZH2) **(Fig. 2D; Fig. S6A-E)** and NEPC markers (NCMA1, CHGA, CEACAM5/6)^21^ **(Fig. 2D; Fig. S5B)** by both B18-94 treatment and BRN2 knockout **(Fig. 2D)** mirroring the transcriptional changes induced by chemically distinct parental compounds B7 and B18 **(Fig. S7C)**. As expected for cells treated with a small molecule inhibitor, discordant genes (Q4) upregulated by B18-94 and not in BRN2^-/-^ cells belong to small molecule metabolism pathways **(Fig. S7B)**. Overall, these data clearly demonstrate that inhibition of BRN2 by B18-94 or genetic knockout affect the same biological pathways involved in cell plasticity and proliferation.

**Figure 2:**
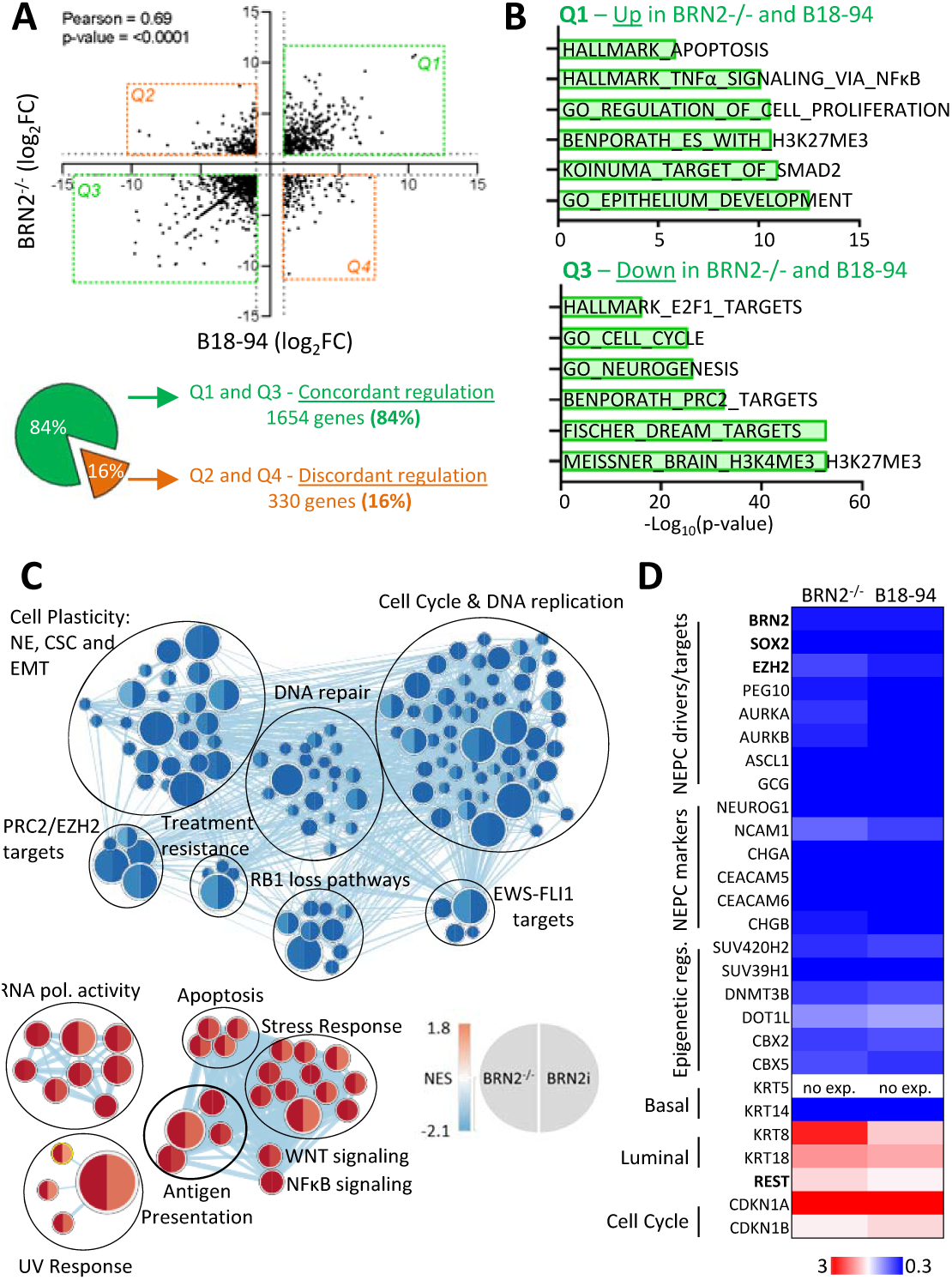
Specificity of BRN2 inhibitor for NEPC models. **(A)** RNA-seq comparing log2(Fold Change vs. Control) between BRN2^-/-^ and B18-94 treated 42D^ENZR^ cells. **(B)** Pathways enriched in each quadrant with concordant and discordant regulation of genes between BRN2^-/-^ and B18-94. **(C)** Biological clustering of GSEA up and downregulated in 42D^ENZR^ cells upon BRN2 inhibition via CRISPR/Cas9 or B18-94 **(D)** Change in expression of specific genes in 42D^ENZR^ cells with Cas9 mediated knockout of BRN2 compared to treatment with B18-94 for 48 hours.

### B18-94 mode of action

Our *in-silico* simulations predicted a 4.7Å shift between the POUs and POU_H_ domains upon binding of B18-94. Therefore, we hypothesized that binding of the small molecule in this pocket closes the distance between the two DBDs, thus, reducing BRN2’s affinity to DNA **(Fig. 1C and 3A)**. To interrogate this hypothesis, we analysed BRN2 binding to the chromatin using fractionation and ChIP-seq. We found that B18-94 drastically reduces the BRN2 bound to chromatin in a dose-dependent manner within 16 hours of treatment, without altering BRN2 nuclear localization. Importantly, overall transcription remained unaffected as RNA polymerase II (POL2RA) binding to the chromatin was unchanged **(Fig. 3A)**. ChIP-seq analysis at the same time-point also confirmed that that B18-94 significantly reduced BRN2 binding to chromatin (**Fig. 3B)**. Moreover, upon analysis of the control (DMSO) ChIP-seq data, we can report for the first time, the BRN2 cistrome in both 42D^ENZR^ and NCI-H660 cells. We observed a strong consistency in biological pathways enriched in the genes with BRN2 binding sites, including neurogenesis and nervous system development **(Fig. S8A-B)**. Within these pathways, specific loci of ASCL1 and SOX2, genes strongly correlating with BRN2 expression in multiple PCa datasets **(Fig. S8C)**, demonstrate a large reduction in BRN2 binding in the promoter region of both genes upon B18-94 treatment **(Fig. 3C)**. This pattern was similarly observed for the NEPC marker PEG10 and BRN2 itself, further supporting the hypothesis for BRN2 auto-regulation^10,22^ **(Fig. S8D)**. Importantly, the lack of BRN2 binding also significantly reduced expression of ASCL1, SOX2 and BRN2 itself in both models by qRT-PCR prior to any observed reduction in BRN2 protein levels **(Fig. S9A)**, confirming that the transcriptional downregulation is due to inhibition of BRN2-chromatin interactions and not a reduction in BRN2 protein levels. These results were further confirmed by an unbiased approach of RNAseq in both 42D^ENZR^ and NCI-H660 cells, as PEG10, ASCL1 and SOX2 were downregulated while enzymes implicated in small molecule metabolism like CYP1A2^23^ and AKR1C2^24^ were upregulated **(Fig. 3D, Table S3-4)**. Similar to the data observed for 42D^ENZR^ cells, NCI-H660 cells also observed significant downregulation in many of NEPC drives/targets/markers that are important for tNEPC progression **(Fig. S9B)**. Biological similarity clustering of gene-set revealed consistent downregulation of cell proliferation, cell plasticity and DNA-repair pathways (a recently reported role of BRN2 in melanoma cells^25^), and upregulation of apoptosis pathways in both models **(Fig. 3D, S7A, S9C)**.

**Figure 3:**
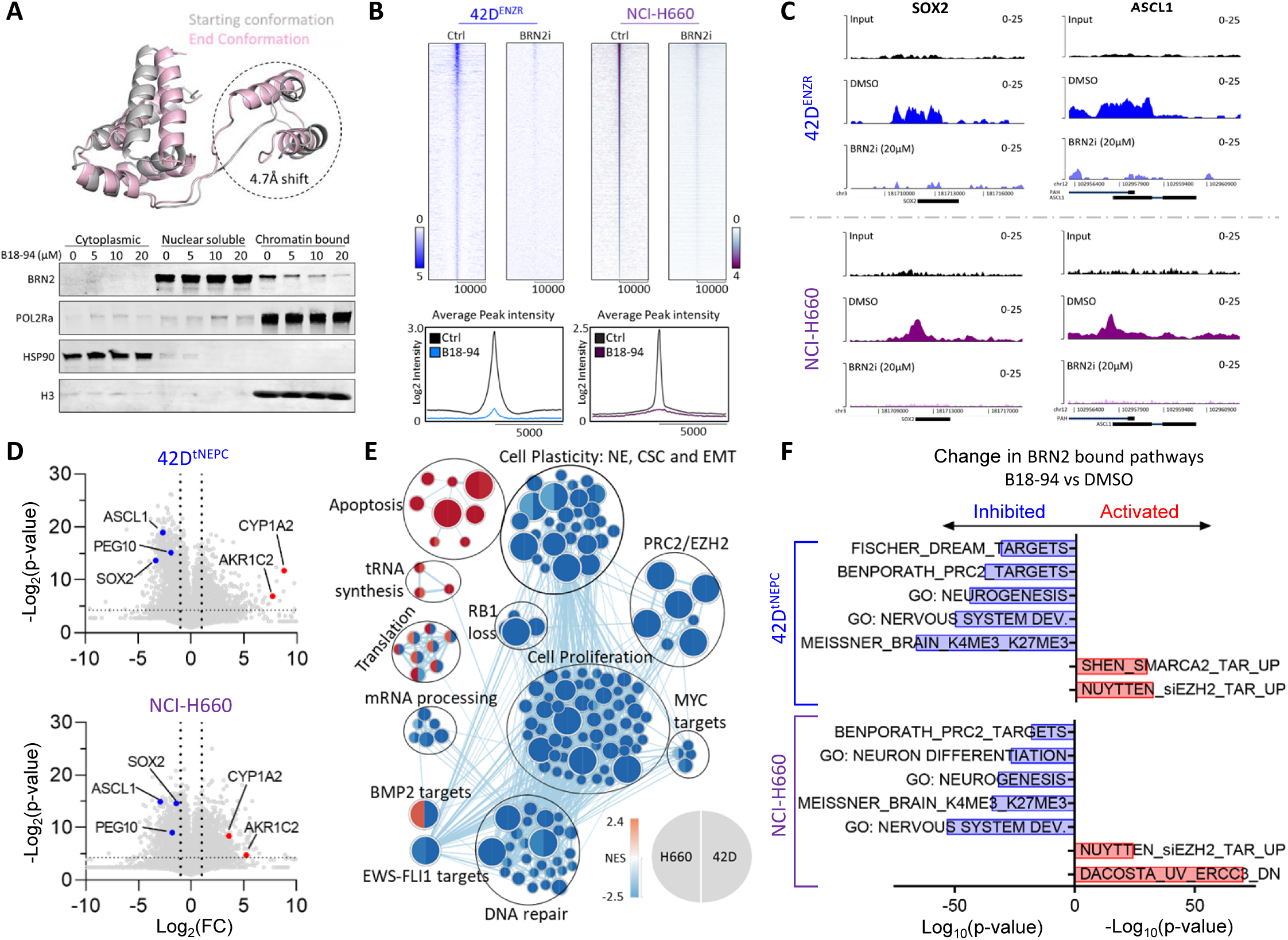
Mechanism of action for BRN2 inhibitor. **(A)** Predicted model on the effect of B18-94 binding to the DBD of BRN2. 42D^ENZR^ cells treated with B18-94 20μM for indicated time points and subject to sub-cellular fractionation. **(B-D)** NCI-H660 and 42D^ENZR^ cells treated with B18-94 20μM Samples were collected and ChIP was performed for BRN2. **(B)** Heat map for BRN2 binding -10000 to +10000 base pairs relative to transcription start sites. **(C)** Annotated tracks from BRN2 ChIP-seq for ASCL1 and SOX2 loci in NCI-H660 and 42D^ENZR^ cells **(D-E)** RNA-seq analysis on NCI-H660 and 42D^ENZR^ cells treated with B18-94. **(D)** Volcano plot showing up-regulated and down-regulated genes **(E)** Biological clustering of upregulated and downregulated pathways upon BRN2 inhibition **(F)** Effect on BRN2 bound pathways in 42D^ENZR^ and NCI-H660 cells observed in **(B)** upon treatment with B18-94 for 48 hours.

Cross-analysis of ChIP-seq and RNA-seq data revealed that the pathways enriched in the BRN2 cistrome like neurogenesis, nervous system development and EZH2/PRC2 targets, were all downregulated upon treatment with B18-94 in both 42D^ENZR^ and NCI-H660 cells **(Fig. 3F)**. In concordance with loss of PRC2 activity, the consistently activated pathway included genes upregulated by siEZH2 treatment in PCa cells^26^. Altogether, these data indicate that B18-94 halts BRN2 recruitment to chromatin and subsequently inhibits BRN2 transcriptional program that support the maintenance of NEPC and cell survival.

### In vivo efficacy of B18-94 inhibitor

The pharmacokinetic and toxicity profile of B18-94 were characterized using the Nu/Nu murine model. We tested B18-94 bioavailability with both intra-peritoneal (IP) and *per os* (PO) routes of administration. Interestingly, both methods converged in their serum concentrations at approximately 8 hours **(Fig. S10A)**. With a calculated half-life of approximately 4.1 hours **(Fig. S10A)**, we proceeded with PO dosing and conducted a multi-dose toxicity study. We discovered that B18-94 is well tolerated even at 200mg/kg without any weight loss or neurotoxicity symptoms (lethargy and repetitive actions) for up to 2 weeks of treatment **(Fig. S10B)**. Taking the 50 and 200 mg/kg doses, we performed an in vivo dose response experiment with NCI-H660 xenografts and found no significant advantage from the higher dose **(Fig. S10C)**. Subsequent studies with larger number of animals per arm clearly demonstrated that 50 mg/kg dose of B18-94 significantly reduced tumor volume of 42D^ENZR^ and NCI-H660 xenograft **(Fig. 4A-B)**. Immunohistochemistry on end-of-study tumors demonstrated clear reduction in BRN2, SOX2 and Ki67 (cell proliferation) as well as an increase in CASP3 (apoptosis) **(Fig. 4A-B, S11)**. Additionally, B18-94 treated tumors had significantly reduced transcript levels of BRN2, SOX2, ASCL1 and PEG10 **(Fig. 4A-B)**. Lastly, we next compared the efficacy of B18-94 to standard of care platinum-based chemotherapy using the NCI-H660 xenografts,. The 50mg/kg dose of B18-94 demonstrated comparable anti-tumor activity to 20 mg/kg dose of carboplatin, however in contrast to carboplatin without the toxicity induced weight loss **(Fig. 4C)**.

**Fig. 4:**
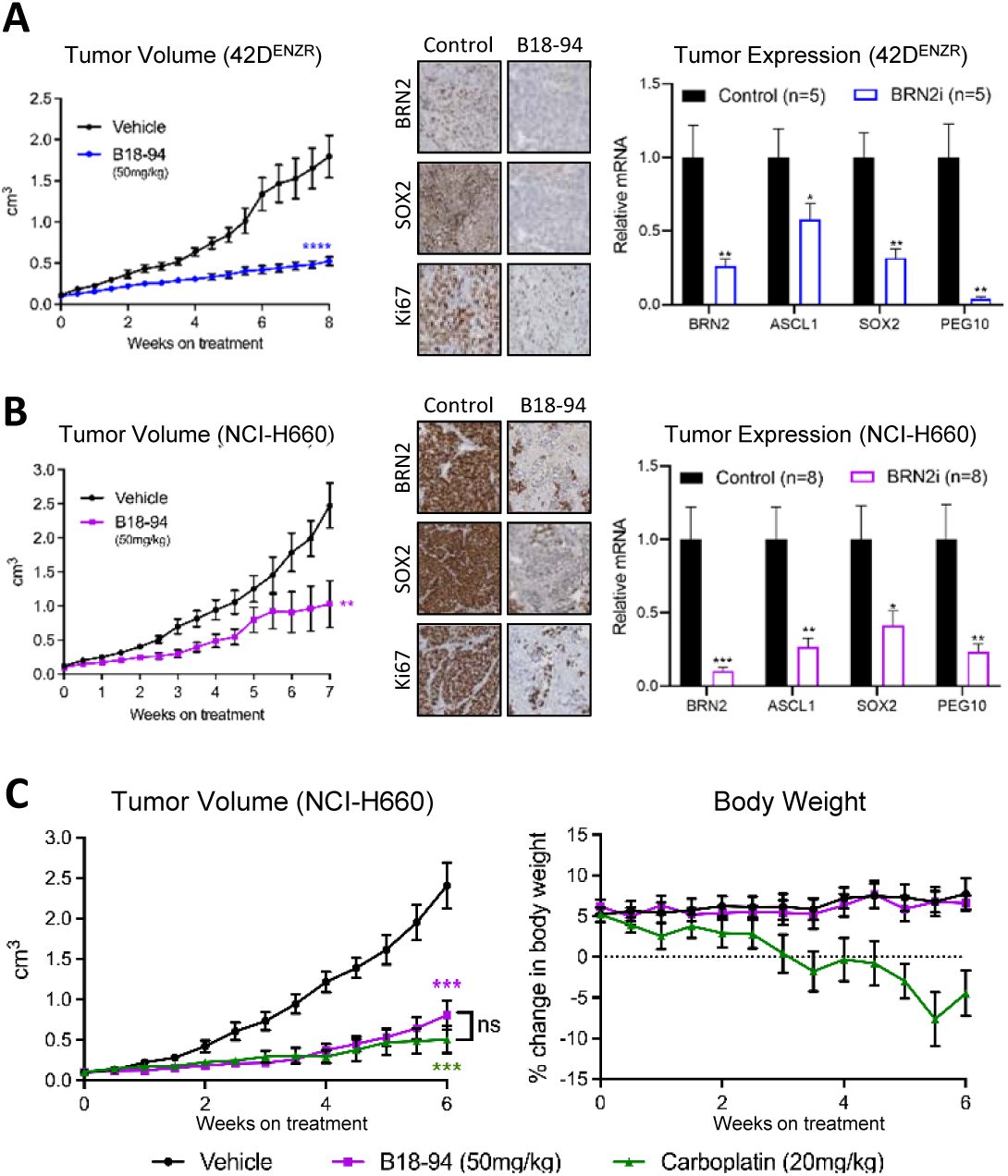
*In vivo* efficacy of BRN2 inhibitor. **(A-B)** Xenograft of 42D^ENZR^ and NCI-H660 injected into Nu/Nu mice. Mice were treated with indicated doses of B18-94 when tumor volume reached 100mm^3^. **(A)** Tumor volume of 42D^ENZR^ xenografts treated with control (n=7) and 50mg/kg of B18-94 (n=10), IHC staining and qRT-PCR for indicated target genes. The results were reported as mean ± SEM; * denotes p<0.05, ** denotes p<0.01, *** denotes p<0.001, **** denotes p<0.0001 **(B)** Tumor volume of NCI-H660 xenografts treated with control (n=8) and 50mg/kg of B18-94 (n=9), IHC staining and qPCR for indicated target genes. The results were reported as mean ± SEM; * denotes p<0.05, ** denotes p<0.01, *** denotes p<0.001 **(C)** Tumor volume and change in body weight of NCI-H660 xenografts treated with vehicle (n=5), carboplatin (n=5) and B18-94 (n=6). The results were reported as mean ± SEM; *** denotes p<0.001

In conclusion, this study characterized the discovery and lead optimization of B18-94, the first-in-class orally bioavailable inhibitor of BRN2. This compound potently inhibits BRN2 recruitment to the chromatin, subsequent transcriptional activity and significantly reduces tumor growth in multiple xenograft models. Overall, these data provide crucial proof-of-concept for targeting BRN2 in NEPC and open avenues for further investigation into other small cell neuroendocrine tumor types.

## Acknowledgements

This work was supported by Prostate Cancer Canada (AZ and RM), Prostate Cancer Foundation (AZ, HB, RM and DT), Michael-Smith Foundation for Health Research (RM), Canadian Institutes of Health Research (AZ and DT), and US-DOD PCRP Idea award (AZ, DT, CM and EC)

## Author Contributions

DT, RM and AZ designed the study. RM performed hit identification and hit-to-lead optimization. DT, SK, SJ and SV did the *in vitro* small-molecule testing. AA conducted x-ray crystallography. DT, SJ and OS conducted *in vivo* studies and processed tumors for IHC and qRT-PCR. DG, JB, SK, HB, CM and EC provided disease expertise and edited manuscript. DT performed all other functional assays and created figures, including performing and analyzing ChIP-seq & RNA-seq. DG analyzed ChIP-seq. DT, RM and AZ wrote the manuscript. All authors had intellectual contribution and approved the manuscript and the data within it.

