## Supplementary figures 1-11 for "Discovery and characterization of a first-in-field transcription factor BRN2 inhibitor for the treatment of neuroendocrine prostate cancer"

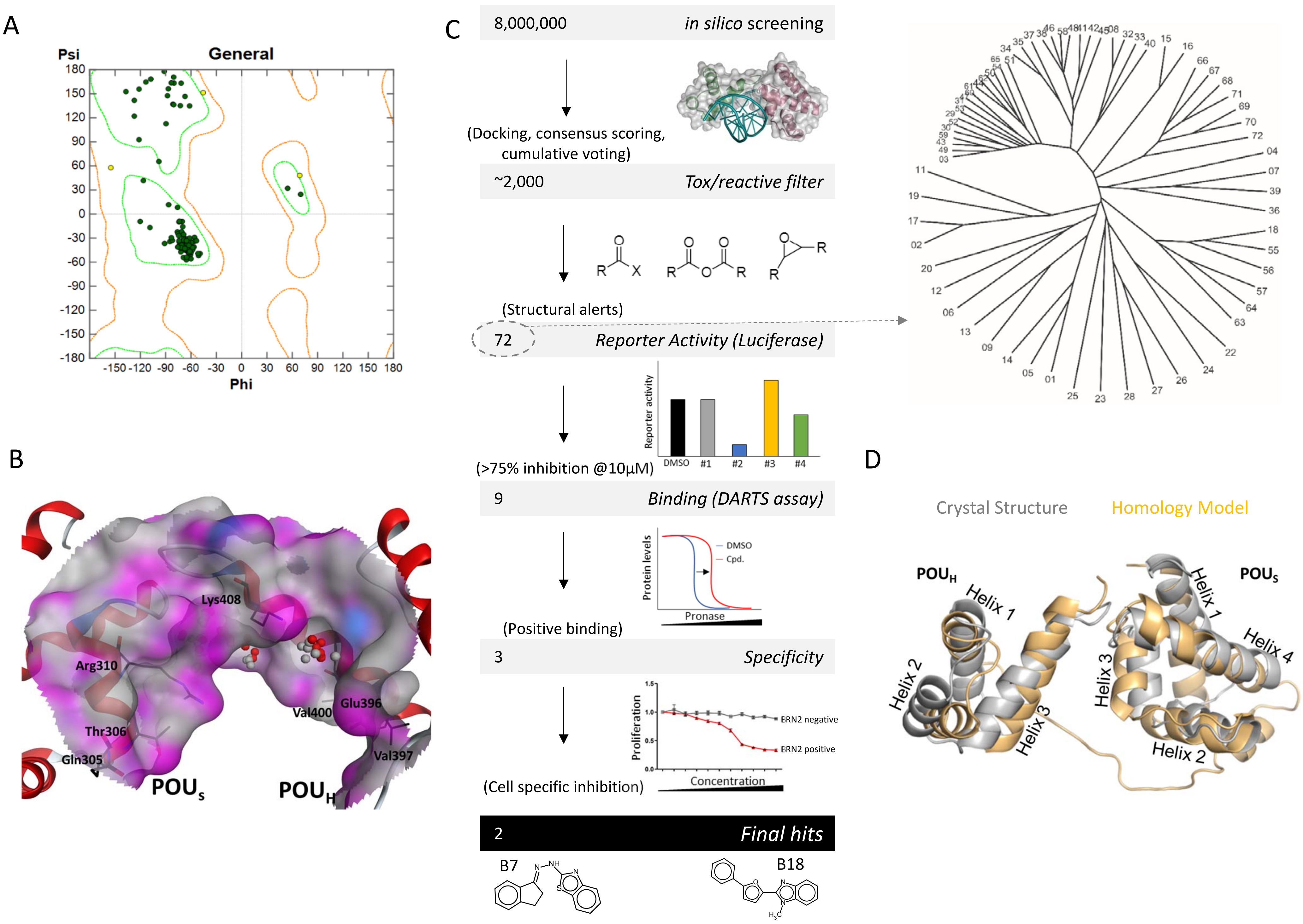
A
Psi
General
Phi
B
Lys408
Arg310
Glu396
Val400
Thr306
Gln305
Val397
$POU_S$
$POU_H$
C
8,000,000
in silico screening
(Docking, consensus scoring, cumulative voting)
~2,000
Tox/reactive filter
(Structural alerts)
72
Reporter Activity (Luciferase)
Reporter activity
DMSO
#1
#2
#3
#4
(>75% inhibition @10μM)
9
Binding (DARTS assay)
Protein levels
DMSO
Cpd.
Pronase
(Positive binding)
3
Specificity
Proliferation
ERN2 negative
ERN2 positive
Concentration
(Cell specific inhibition)
2
Final hits
B7
B18
D
Crystal Structure
Homology Model
$POU_H$
$POU_S$
Helix 1
Helix 2
Helix 3
Helix 4
Helix 1
Helix 3
Helix 2

Supplementary S2 - related to Figure 1

A

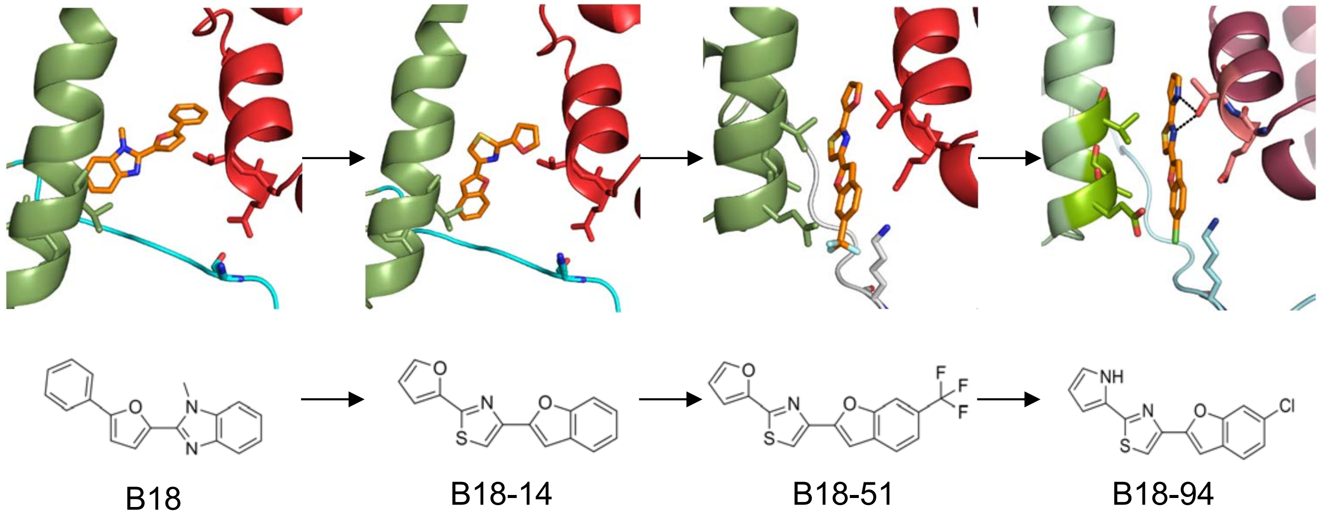


B

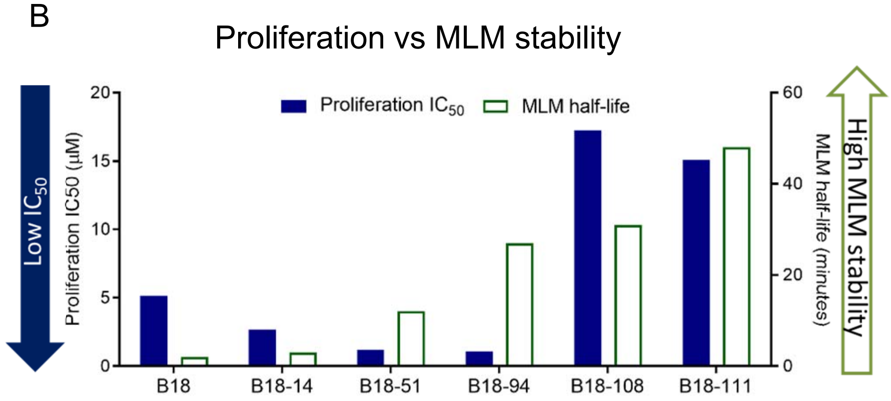


C

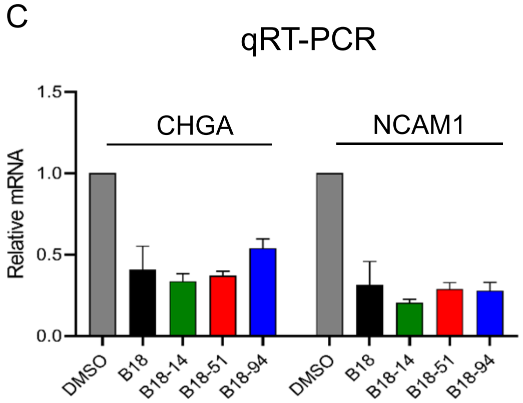

Supplementary S3 - related to Figure 1

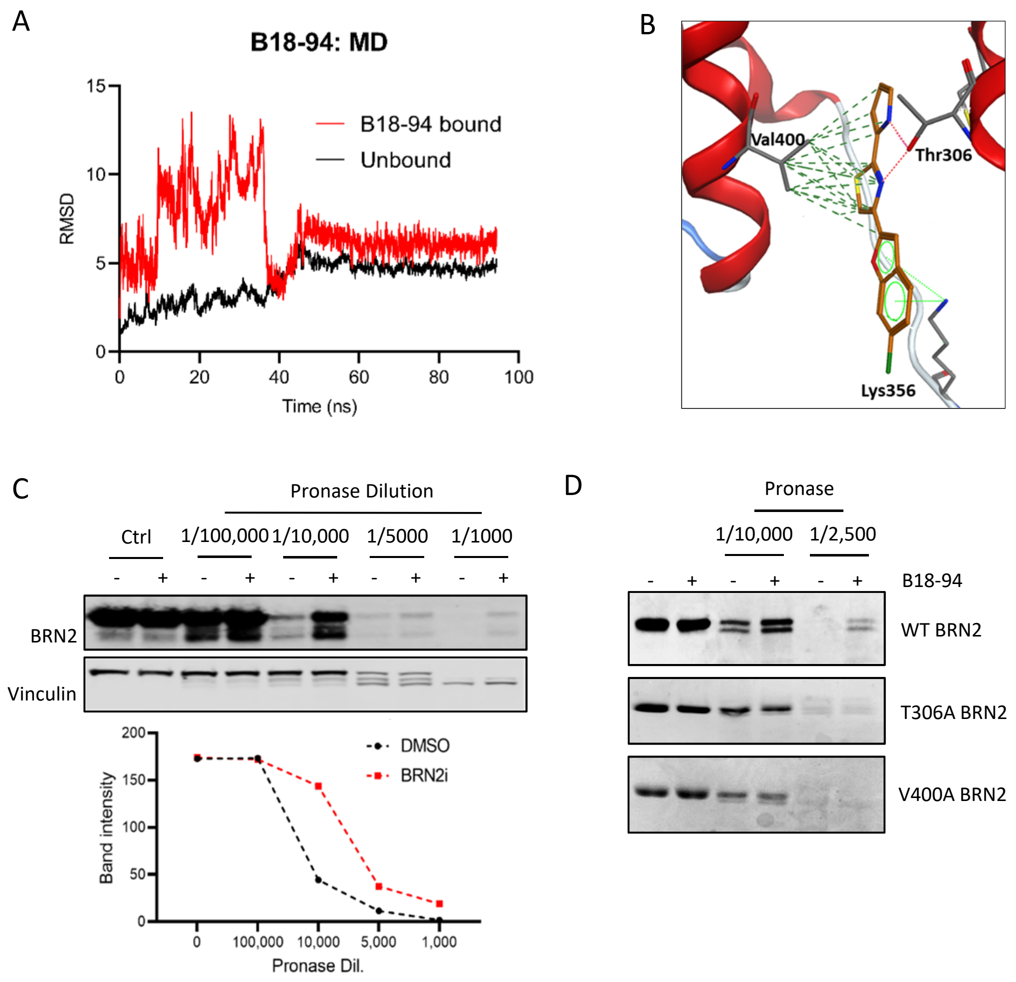

### Supplementary S4 – Related to specificity data

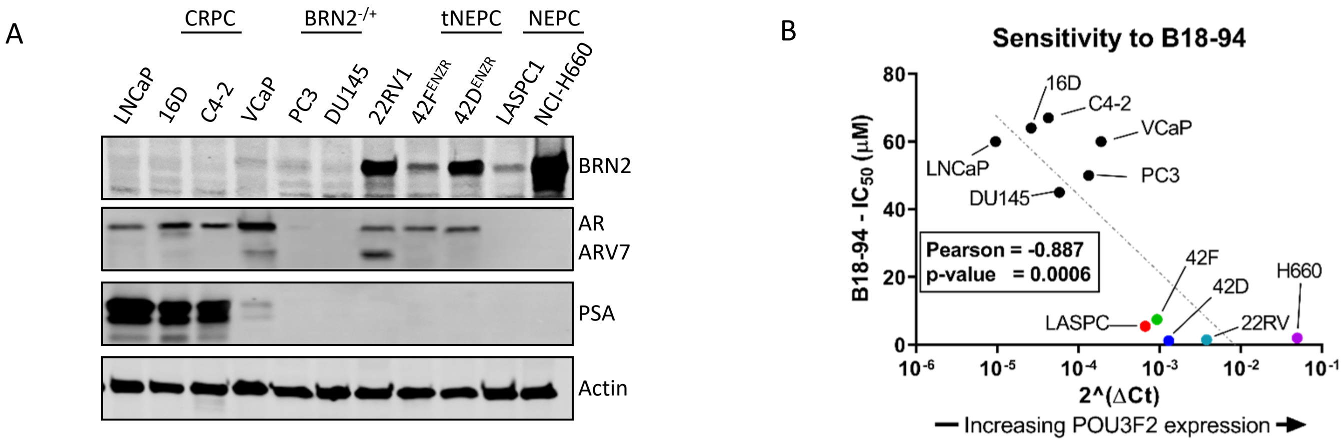


C

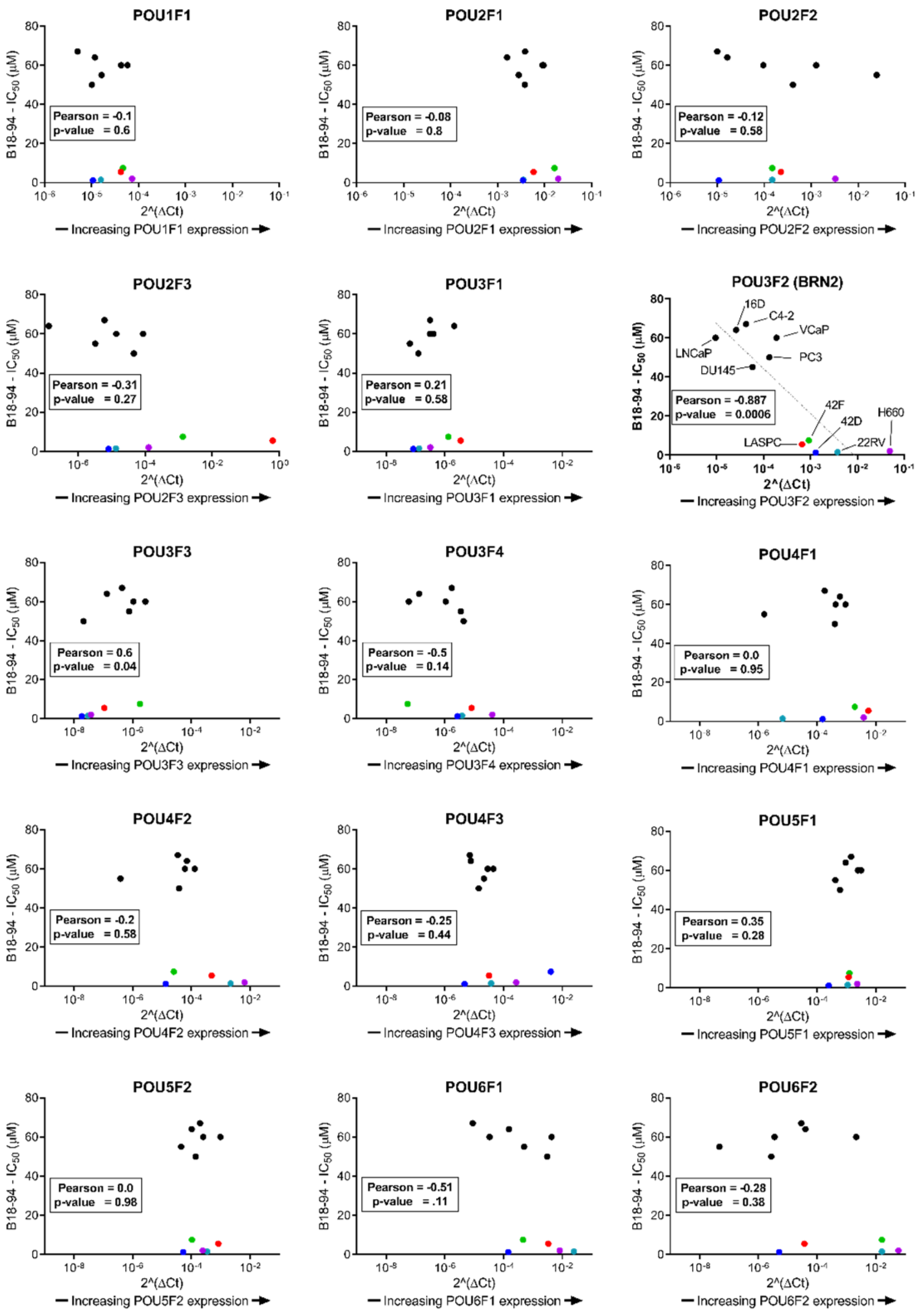


| Gene | Pearson | p-value |
|---|---|---|
| POU1F1 | -0.1 | 0.6 |
| POU2F1 | -0.08 | 0.8 |
| POU2F2 | -0.12 | 0.58 |
| POU2F3 | -0.31 | 0.27 |
| POU3F1 | +0.21 | 0.58 |
| ***POU3F2*** | ***-0.887*** | ***0.0006*** |
| ***POU3F3*** | ***+0.6*** | ***0.04*** |
| POU3F4 | -0.5 | 0.14 |
| POU4F1 | 0 | 0.95 |
| POU4F2 | -0.2 | 0.58 |
| POU4F3 | -0.25 | 0.44 |
| POU5F1 | +0.35 | 0.28 |
| POU5F2 | 0 | 0.98 |
| POU6F1 | -0.51 | 0.11 |
| POU6F2 | -0.29 | 0.38 |

Supplementary S5 – Related to specificity data

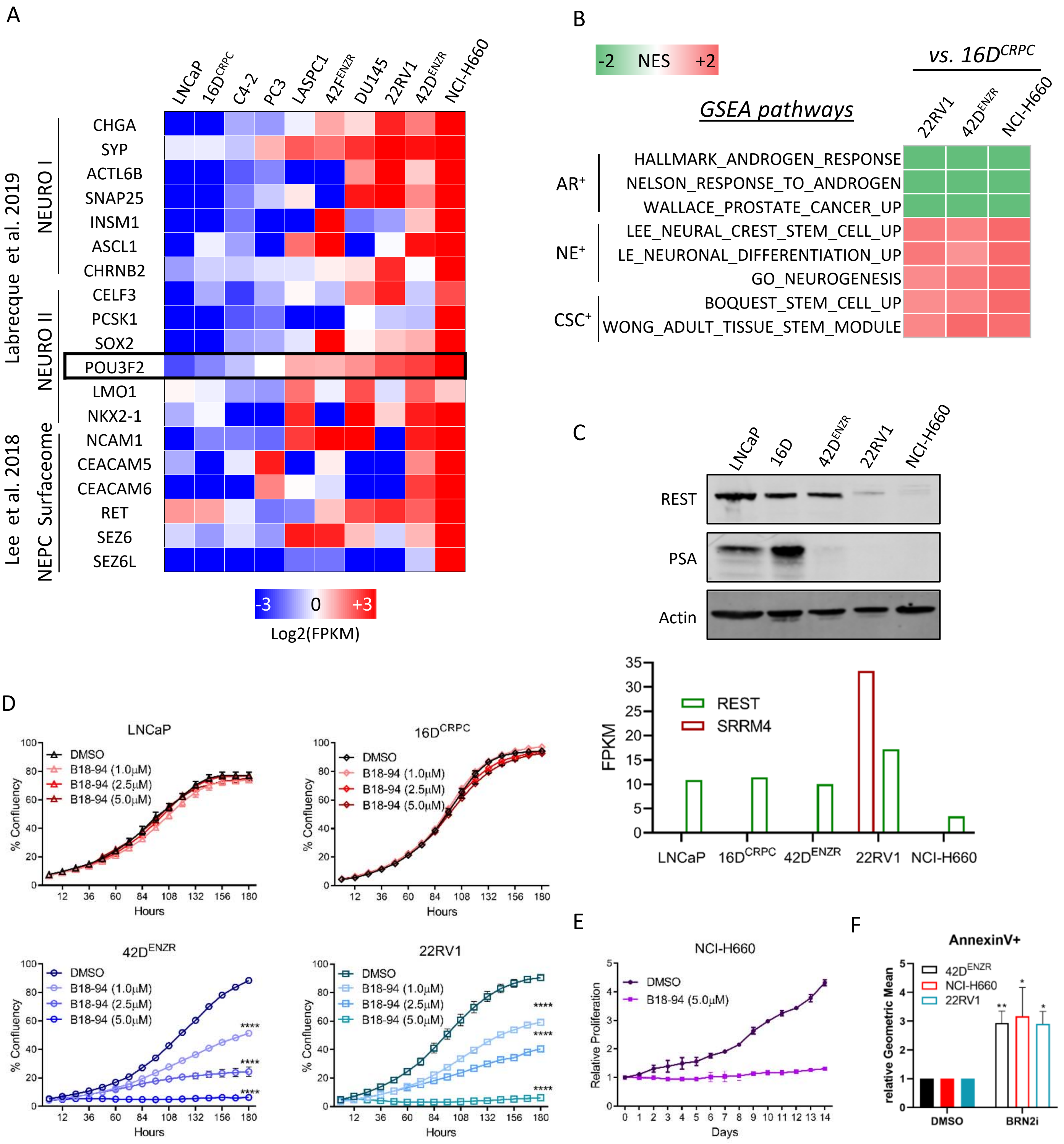

### Supplementary S6 – related to specificity

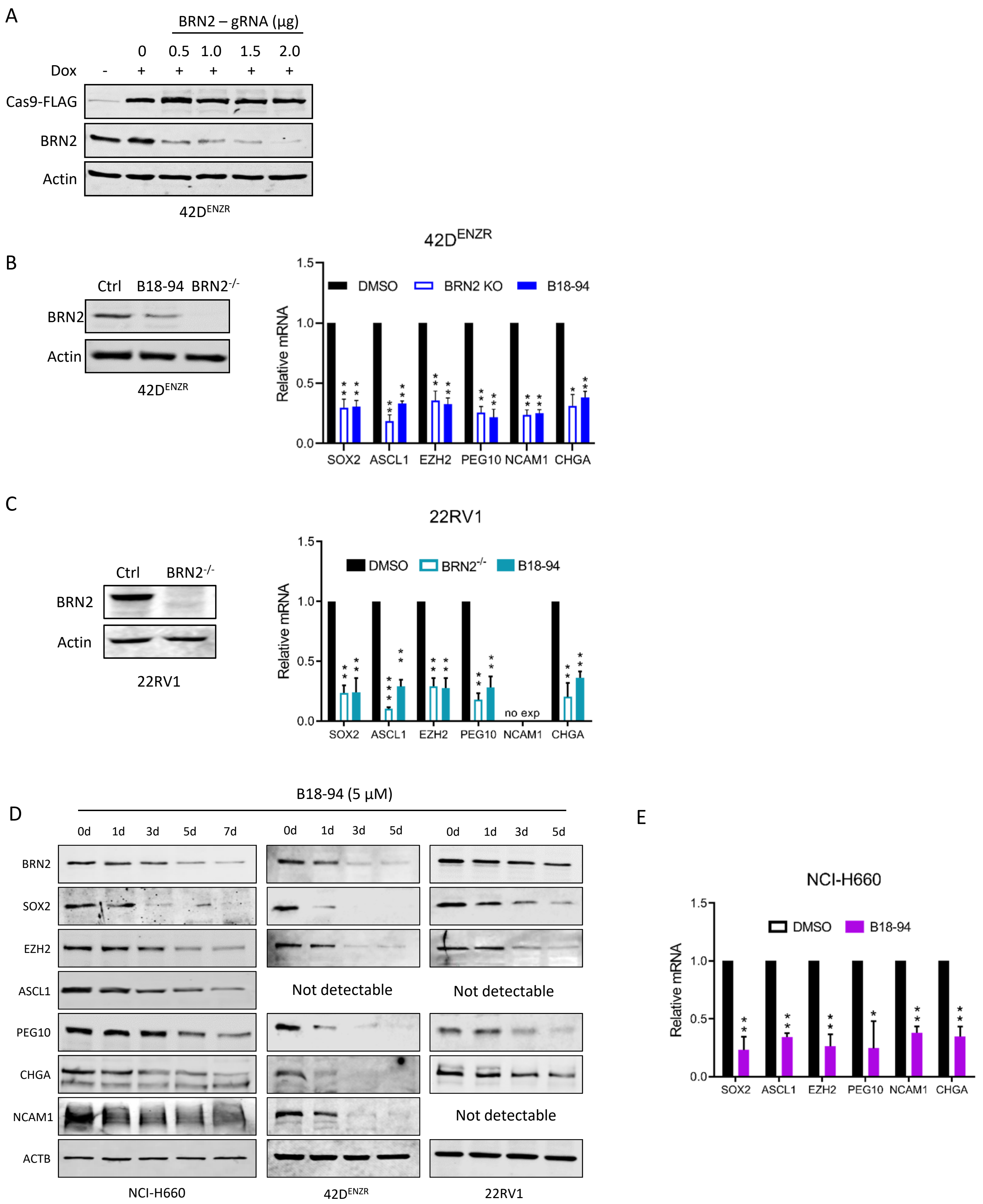

Supplementary S7 – related to specificity

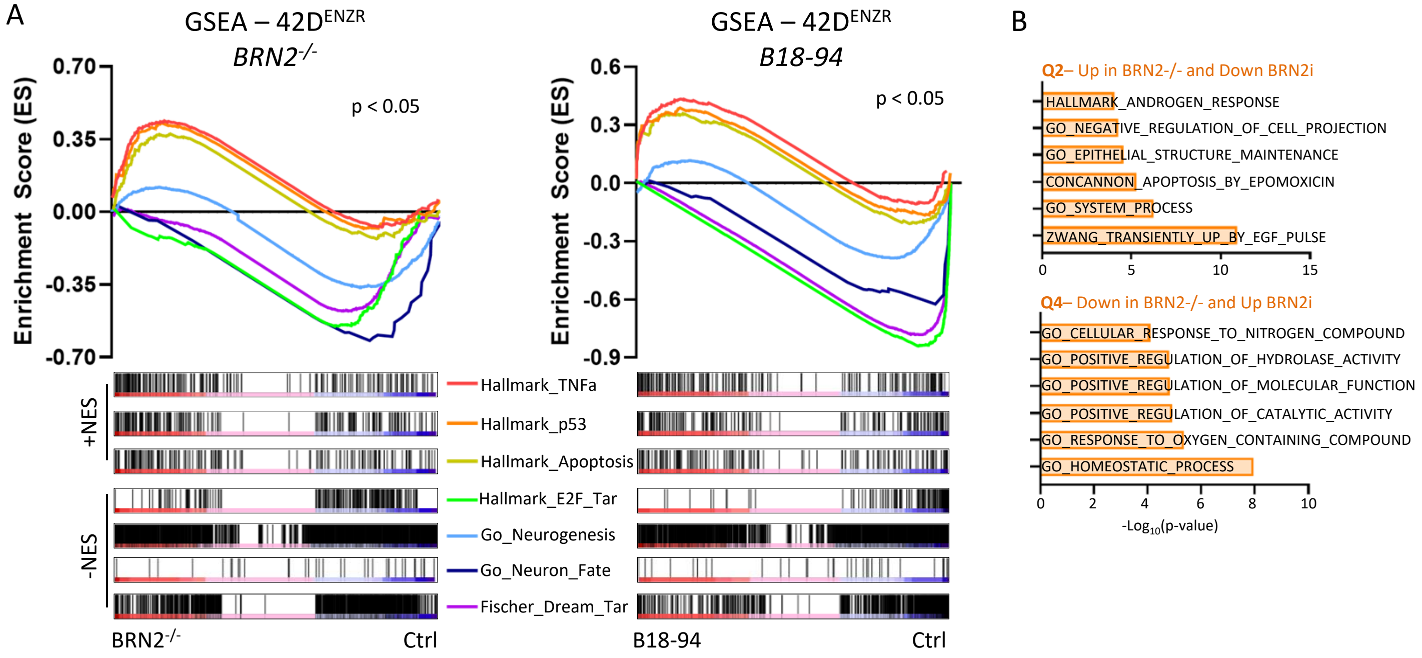


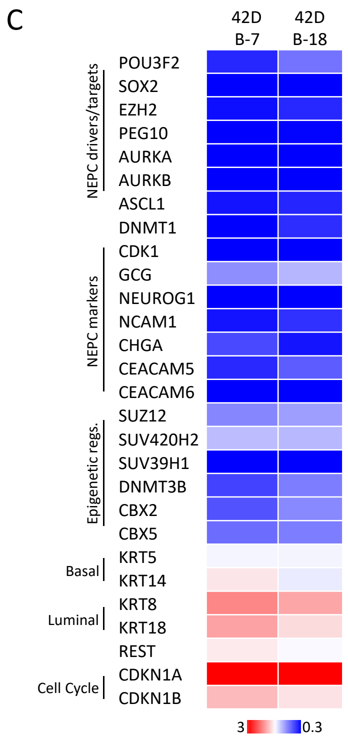

### Supplementary S8 – related to MOA

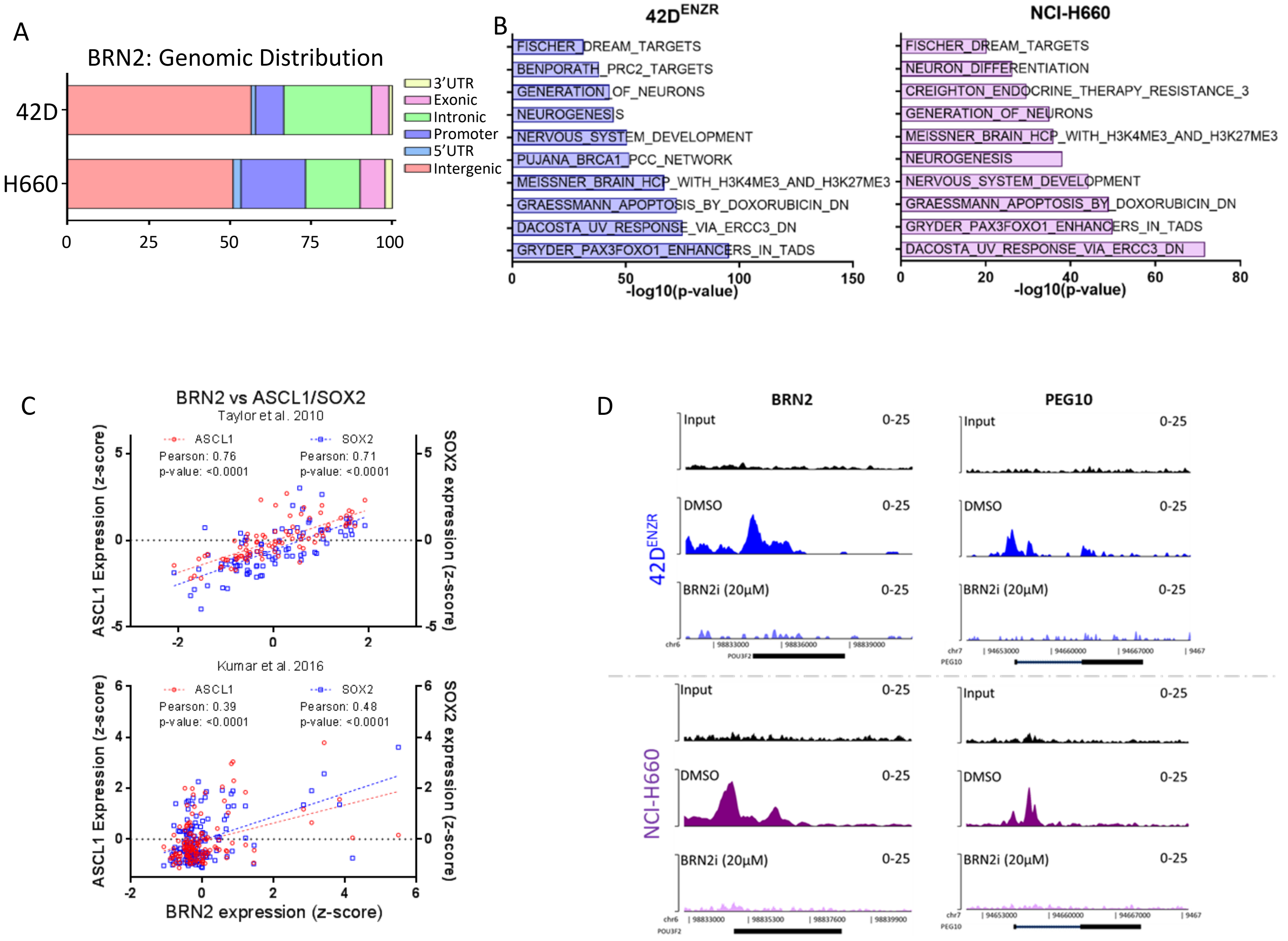

### Supplementary S9 – related to MOA

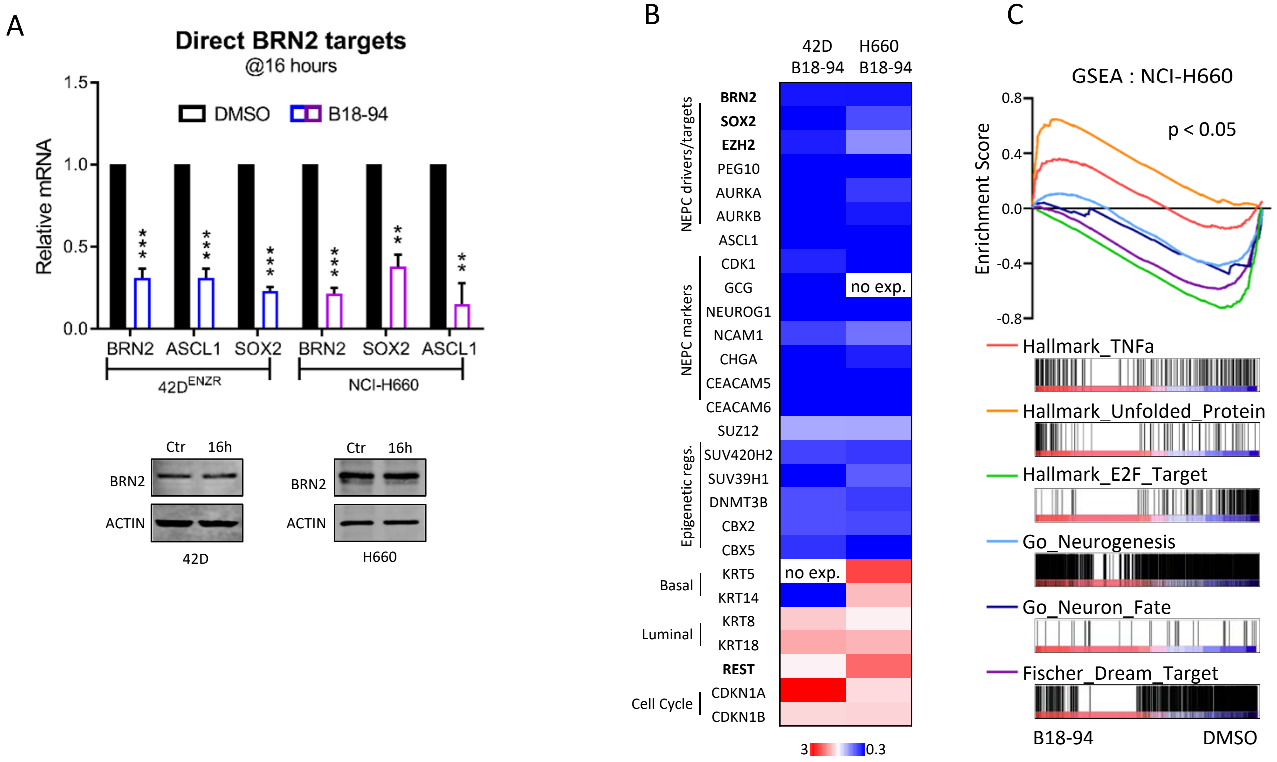

### Supplementary D10: in vivo data

A

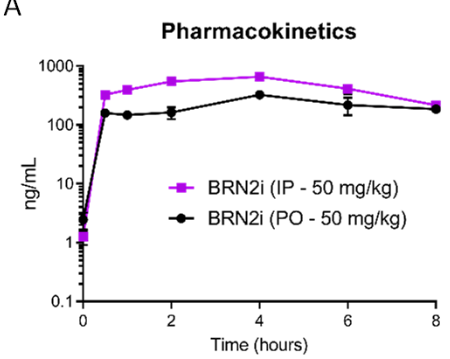


| pK parameter | Unit | Estimate |
|---|---|---|
| $T_{1/2}$ | hours | 4.1 |
| $T_{Max}$ | hours | 4 |
| $C_{Max}$ | ng/mL | 325 |

B

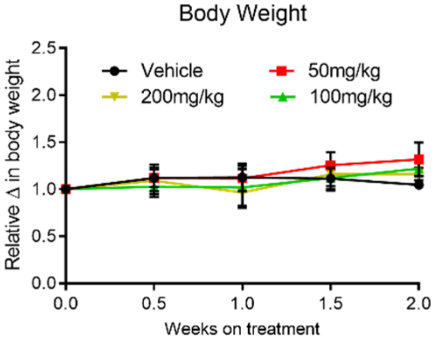


C

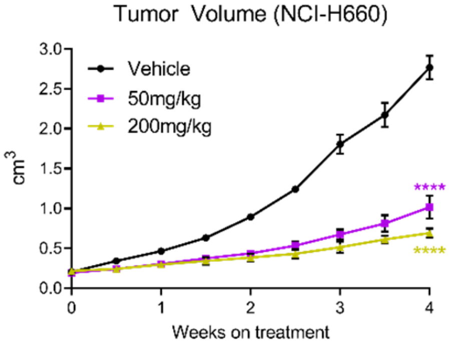

Supplementary D11: in vivo data

A

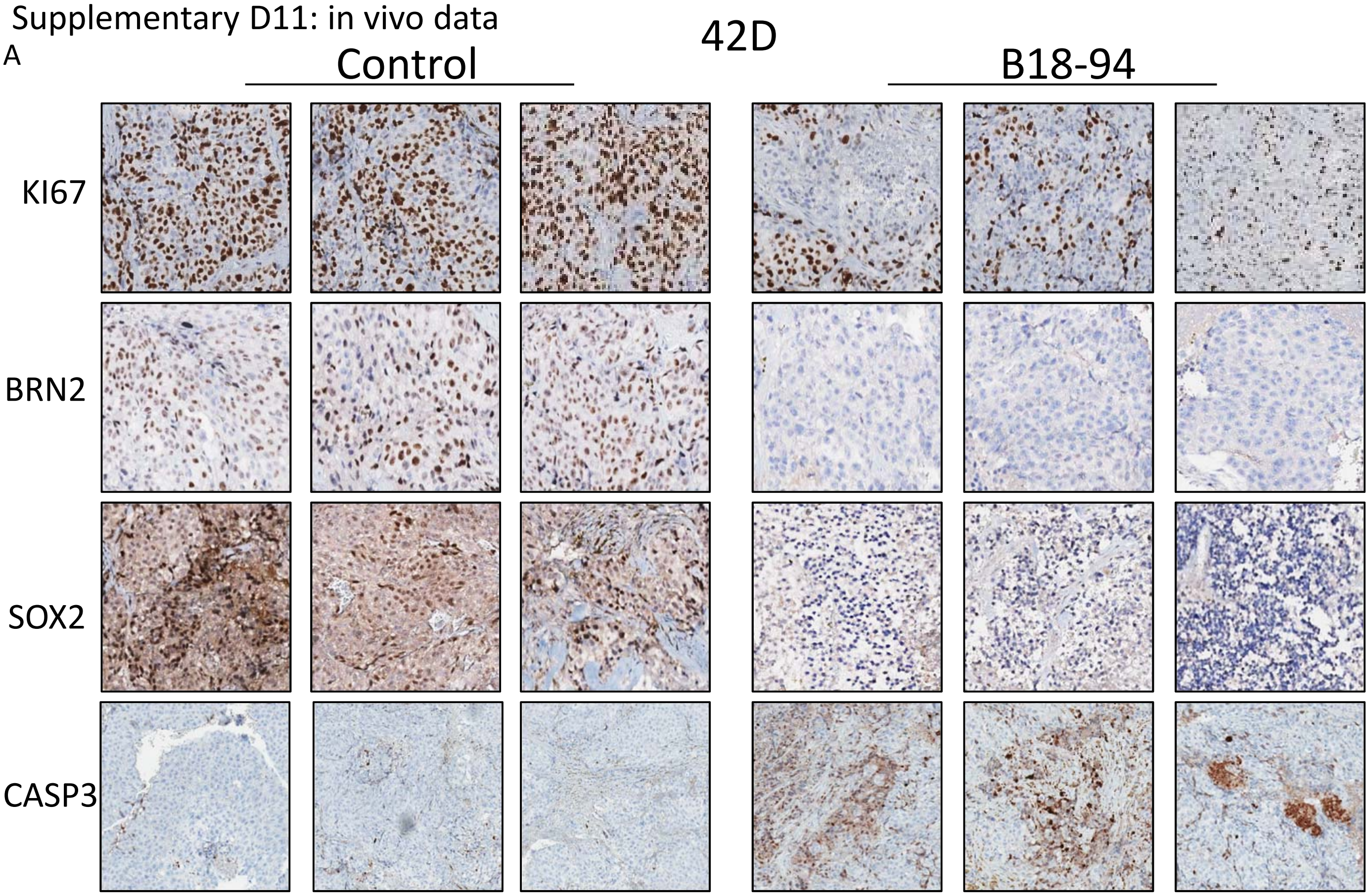


B

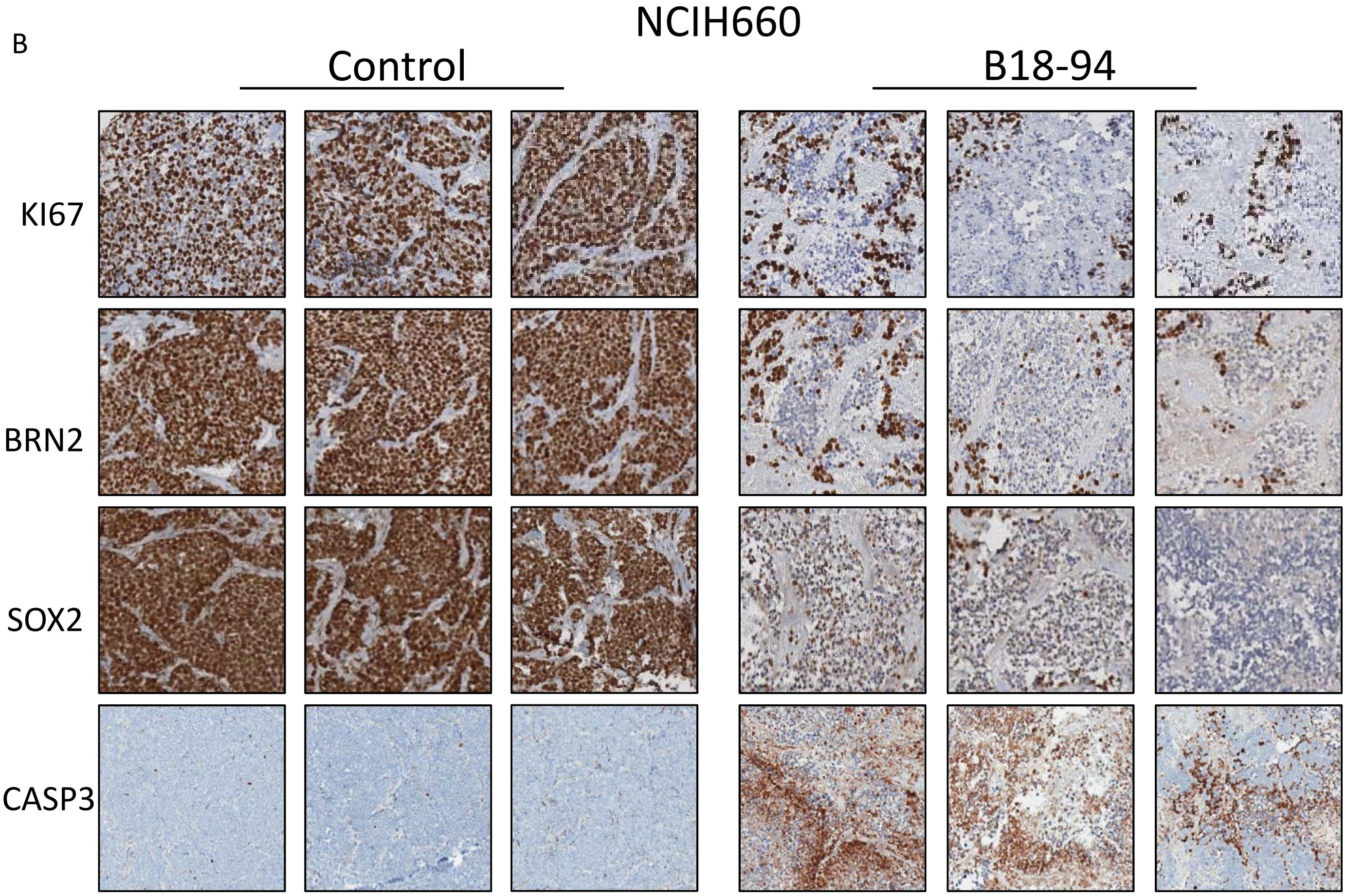
