## Supplementary figure legends for "Discovery and characterization of a first-in-field transcription factor BRN2 inhibitor for the treatment of neuroendocrine prostate cancer"

**Figure S1: Identification of BRN2 inhibitory chemical scaffolds B7 and B18.**

**(A)** Ramachandran plot showing the quality of the BRN2 structure developed using homology modeling approach. Green and yellow dots represent the amino acids that fall into the "allowed" and "generously allowed" regions, respectively. **(B)** Predicted solvent-exposed binding site on the human BRN2 X-ray structure. The dummy atoms enveloped by residues indicate potentially contributes to hydrophobic and hydrogen bond interactions in the region between POUH and POUS domains. The grey and red spheres indicate hydrophobic and polar areas, respectively. **(C)** *Small molecule selection pipeline:* We began with an in-silico screen of ZINC database against the BRN2-DBD pocket. Top 2000 molecules were passed to a toxic moiety filter and seventy-two structurally diverse (as shown in unrooted similarity tree) compounds were selected for further investigation. Luciferase reporter assay: $42D^{ENZR}$ stably expressing BRN2 reporter treated with 10uM BRN2 inhibitors and luciferase activity measured. DARTS assay: Cell lysate was incubated with BRN2 inhibitors in the presence or absence of Pronase. Cell proliferation: BRN2 positive $42D^{ENZR}$ and BRN2 negative $16D^{CRPC}$ were treated with increasing concentrations of BRN2 inhibitors for 72 hours and cell proliferation was performed. **(D)** Overlay of BRN2-DBD crystal structure with the in-silico derived BRN2 homology model.

**Figure S2: Lead optimization of scaffold B18.**

**A)** Docking of BRN2-DBD with chemical structures of parental compound B18 and its derivatives B18-14, B18-51 and B18-94. **B)** Mouse liver microsomal (MLM) stability vs. IC50 $42D^{ENZR}$ cells of B18 derivatives. C) qRT-PCR for BRN2 target genes and NEPC markers CHGA and NCAM1 in $42D^{ENZR}$ treated with 10μM of B18, B18-14, B18-51 and B18-94 for 48 hours.

**Figure S3: B18-94 binding to BRN2 DNA-binding domain.**

**A)** RMSD curves demonstrating the individual stabilities of BRN2-DBD along or bound to B18-94 during MD simulations. **B)** Predicted interactions between specific residues of BRN2 with B18-94. **C)** DARTS assay: Cell lysate incubated with DMSO or B18-94 exposed to increasing dilutions of Pronase stock at 12.5μg/mL, followed by Western Blot for BRN2 and Vinculin. Densitometry readings from ImageStudio are graphed below. **D)** DARTS assay of B18-94 binding against purified $BRN2^{WT}$, $BRN2T^{306A}$ and $BRN2^{V400E}$.

**Figure S4: Specificity of B18-94.**

**A)** Western Blot for BRN2, AR, PSA and Actin in a panel of prostate cancer cell lines. **B)** Scatter plot of $IC_{50}$ of prostate cancer models vs. their respective mRNA expression of BRN2. **C)** Scatter plot of $IC_{50}$ of prostate cancer models vs. their respective mRNA expression of POU family members. **D)** Table containing correlation and p-value of IC50 vs POU family gene expression across prostate cancer models.

**Figure S5: B18-94 inhibits prostate cancer models with NEPC phenotype.**

**A)** Heatmap of FPKM values of RNA-seq data of prostate cancer models for indicated genes. **B)** Heatmap of GSEA NES values comparing 22RV1, 42D and NCI-H660 to $16D^{CRPC}$ cells. **C)** Top: Protein expression of REST, PSA and Actin in indicated PCa cell lines. Bottom: mRNA (FPKM) values of REST and SRRM4 in indicated PCa cell lines. **D-E)** Dose response of cell proliferation over time for LNCaP, $16D^{CRPC}$, NCI-H660, $42D^{ENZR}$ and 22RV1 cells treated with B18-94. **F)** Annexin V staining for NCI-H660, $42D^{ENZR}$ and 22RV1 cells treated with B18-94. NCI-H660 cells were treated with 10µM B18-94 for 72 hours while $42D^{ENZR}$ and 22RV1 cells were treated with 5µM for 48 hours.

**Figure S6: B18-94 and BRN2 knockout reduce NEPC target gene expression**

**A)** $42D^{ENZR}$ cells stable expressing rtA-eSpCas9-FLAG are treated Doxycycline and with increased concentration of gRNA targeting Exon 1 of BRN2. **B)** Left: WB for BRN2 and Actin in $42D^{ENZR}$ cells treated with 5µM B18-94 or 2µg of gBRN2. Right: qRT-PCR comparing relative expression of BRN2 target genes upon treatment with B18-94 or BRN2 knockout via Cas9/gRNA. The results were reported as mean ± SD; * denotes p<0.05, ** denotes p<0.01, *** denotes p<0.001 **C)** Left: WB validating BRN2 knockout in 22RV1 cells. Right: qRT-PCR comparing relative expression of BRN2 target genes upon treatment with B18-94 or BRN2 knockout via Cas9/gRNA. The results were reported as mean ± SD; * denotes p<0.05, ** denotes p<0.01, *** denotes p<0.001 **D)** WB for indicated genes in time-course of NCI-H660, $42D^{ENZR}$ and 22RV1 cells treated with 5µM of B18-94.

**Figure S7: B18-94 broadly phenocopies BRN2 knockout.**

**A)** Individual GSEA pathways for Apoptosis, Neuronal and Cell Cycle pathways upon inhibition of BRN2 by B18-94 or Cas9 mediated knockout. **B)** Pathways enriched in Q2 and Q4 containing discordantly regulated genes between $BRN2^{-/-}$ and B18-94.

**Figure S8: BRN2 cistrome is enriched for neuronal and cell cycle pathways.**

**A)** Genomic distribution of BRN2 binding sites relative to gene bodies within the chromatin. **B)** Pathway enriched in genes mapped to BRN2 binding sites in the genome in tNEPC $42D^{ENZR}$ and de novo NEPC NCI-H660 model. **C)** Correlation of BRN2 vs. ASCL1 and BRN2 vs. SOX2 in two metastatic prostate cancer datasets. **D)** Annotated tracks from BRN2 ChIP-seq for BRN2 and PEG10 loci in NCI-H660 and $42D^{ENZR}$ cells.

**Figure S9: B18-94 treatment suppresses proliferative and NEPC phenotypes.**

**A)** qRT-PCR analysis for BRN2, SOX2 and ASCL1 in NCI-H660 and $42D^{ENZR}$ cells treated with 20µM of B18-94 for 16 hours. The results were reported as mean ± SD; * denotes p<0.05, ** denotes p<0.01, *** denotes p<0.001 **B)** Heatmap of normalized RNA-seq data (fold change vs. DMSO) of NCI-H660 and $42D^{ENZR}$ cells treated with B18-94. NCI-H660 cells were treated with 10µM B18-94 for 72 hours while $42D^{ENZR}$ cells were treated with 5µM for 48 hours. **C)** Individual GSEA pathways from RNA-seq for Apoptosis, Neuronal and Cell Cycle pathways upon inhibition of BRN2 by B18-94 in NCI-H660 cells.

**Figure S10: Pharmacokinetics and dosing of B18-94.**

**A)** *Left:* Oral and Intraperitoneal dosing of 50mg/kg of B18-94 in Nu/Nu mice (n=5 per arm). Blood was collected via tail bleed and tested for presence of B18-94 by LCMS. *Right:* Half-life, Tmax and Cmax for oral dosing of B18-94. **B)** Change in body-weight of mice orally treated with different doses of B18-94 over 2 weeks. **C)** Xenograft of NCI-H660 injected into NOD/SCID mice. Mice were treated with indicated doses of B-18-94 when tumor volume reached 200mm$^3$. Pilot study, n=4 per arm.

**Figure S11: IHC expression of BRN2 target genes and proliferation/apoptosis markers.**

**A-B)** Representative IHC staining for BRN2, SOX2, Ki67 and CASP3 in 42D$^{ENZR}$ and NCI-H660 xenografts treated with vehicle control or 50mg/kg of B18-94 from Figure 4A-B.
