## Supplementary material for "Discovery and characterization of a first-in-field transcription factor BRN2 inhibitor for the treatment of neuroendocrine prostate cancer": Materials and Methods

**Homology modelling of BRN2 protein structure:**

Based on the sequence homology and crystal structure of other POU-domain proteins namely Pit1 (PDB code – 1AU7, 2.30Å), Oct1 (PDB code - 1CQT, 3.20 Å) and BRN5 (PDB code - 3D1N, 2.51 Å), a BRN2 structure was generated using Modeller^1^.  The flexible loop that connects POUH and POUs domains was further refined by *Loop Modeler* module in MOE v2015^2^. The quality of the developed BRN2 structure was assessed using Ramachandran plot and 100-ns molecular dynamic (MD) simulations using AMBER^3^. The system was simulated in the NPT ensemble using the Nose-Hoover Langevin piston method to maintain the pressure at 1 atm and the Langevin thermostat was employed to maintain the temperature at 300 K. All MD trajectories were visualized and analyzed using VMD with Tcl scripts.

**Protein, ligand preparation and virtual screening:**

To prepare the *in silico* modeled and crystallographic structures of BRN2, bond orders were adjusted and hydrogen atoms were added. The structures were energy-minimized using the OPLS force field as implemented by Maestro^4^. The ligand-binding region is defined by a 12 Å box centered on the residue Thr306. No van der Waals scaling factors were applied; the default settings were used for all other adjustable parameters.

To prepare ligands, the ZINC database^5^ containing 4 million molecules was used for virtual screening. The compounds were imported into a molecular database using MOE. Hydrogen atoms were added after these structures were “washed” (a procedure including salt disconnection, removal of minor components, deprotonation of strong acids, and protonation of strong bases). The following energy minimization was performed with the MMFF94x force field, as implemented by the MOE, and the optimized structures were exported into the Maestro suite in SD file format.

Finally, 4 million small-molecules were docked into the defined active site grid using Glide^6^ and AutoDock ^7^ programs. Next, RMSD values were calculated between the docking poses generated by Glide and AutoDock to identify the most consistent binding orientation of the compounds. Only molecules with poses having RMSD values below 2.0 Å were selected for further analysis. Furthermore, the selected ligands were subjected to additional scoring functions implemented in MOE. Finally, 2,000 compounds that consistently demonstrated high predicted binding affinities were selected for visual inspection to filter out compounds containing PAINS moieties, reactive or toxic groups. Based on chemical diversity and availability, 72 chemicals were purchased from commercial vendors such as Enamine, VITAS-M and ChemBridge.

**Molecular dynamics simulations on B18-94-BRN2 complex:**

Molecular dynamics (MD) simulations of B18-94 was performed starting from its docked pose in the DBD region of hBRN2 as predicted by Glide. The protein-ligand system was set up as described above. The simulation was performed with the CUDA accelerated Amber^3^. AR LBD force field parameters were obtained from the ff14SB force field and the ligand parameters came from generalized amber force field with charges derived from a RESP fit using an HF/6-31G* electrostatic potential calculated using the Gaussian 09 program. MD simulations were carried out within AMBER on WestGrid facilities from Compute Calculation Canada (https.//www.westgrid.ca).

**Protein Purification:**

Residues 264-415 of BRN2 were cloned into a pSY7 plasmid (gift from Dr. Robert Robinson Lab) and transformed into BL21 expression cells. 50 mL of overnight culture of DBD and incubated at 37°C was added to 950 mL of Terrific broth-Ampicillin with shaking at 200 rpm till OD^600^ = 1. Culture temperature was reduced to 16°C and protein expression was induced with 0.2mM IPTG overnight. Cells were harvested by centrifuging in a Sorval SLA 16.50 rotor at 9000 rpm for 10 minutes. Pellets were re-suspended in lysis buffer and frozen at -30°C.

Protein extraction was achieved by thawing frozen cells and sonicating in the presence of cOmplete protease inhibitor cocktail (*Roche - 11697498001*). Cell debris was removed by centrifugation at 12000 rpm for 45 minutes in a Sorval 25.50 rotor. His-tagged BRN2 was purified via Ni-NTA column purification and incubated overnight with 300µL of precision protease enzyme (Purified in-house) to cleave off the MBP-His tag. This digest was run through an Amylose column which bound MBP-His and allowed unbound and cleaved BRN2-DBD to be collected in flow-through. The final purification step involved passage through a superdex 200 16/60 column. Fractionated BRN2-DBD was utilized for crystallization and binding assays.

Buffers listed below:

Ni-NTA lysis/wash buffer - 50 mM Tris pH 7.6, 500 mM NaCl, 1 mM β-Me and 10 mM imidazole.

Ni-NTA Elution buffer - 50 mM Tris pH 7.6, 500 mM NaCl, 1 mM β-Me and 250 mM imidazole.

Amylose wash buffer - 50 mM Tris pH 7.6, 500 mM NaCl, 1mM β-Me.

Amylose Elution buffer - 50 mM Tris pH 7.6, 500 mM NaCl, 1mM β-Me and 20 mM maltose.

Superdex running buffer - 50 mM Tris pH 7.6, 200 mM NaCl, 1mM β-Me.

**X-Ray Crystallography:**

Purified DBD protein was concentrated to a final concentration of 3 mg/mL. DBD-DNA complex was formed at a protein:DNA molar ratio of 1:1.5 and crystallized using sitting drop vapour diffusion method. Crystals of DBD-MORE appeared within five minutes and grew to dimensions of 0.3 by 0.5 by 0.7mm over one week. Crystallization solution was 20% PEG 3350 and 0.1M Bis Tris pH 5.5. The crystallization drop was composed of 1µL protein complex and 1µL of crystallization solution. Crystals of DBD-MORE were harvested and cryoprotected with crystallization buffer supplemented with 30% glycerol before flash freezing in liquid Nitrogen.

Data was collected on beamline 9-2 at the Stanford Synchrotron Radiation Lightsource with diffraction at 100K and wavelength 0.998 nm. Data processing to 1.95Å was completed using iMOSFLM ^8^ and scaled in SCALA ^8^. Phasing was achieved via molecular replacement using a CHAINSAW-modified ^9^ model of Oct-6 POU domain bound to DNA (PDB ID: 2XSD) in PHASER ^10^. The final model was built through repeated cycles of PHENIX REFINE ^11^ and manual building in COOT ^12^. Final structure quality was analyzed with SFCHECK and PROCHECK ^8^. The coordinates will be deposited in PDB.

*MORE DNA sequence:* The double stranded MORE DNA sequence GTATGCAAATAAG and its complementary strand CTTATTTGCATAC were obtained from Integrated DNA Technologies (Coralville, USA)

**Cell Culture:**

LNCaP, C4-2, 22RV1, 16D^CRPC^ 42D^ENZR^, 42F^ENZR^ and PC3 cells were cultured in RPMI *(Thermo -11875176)* supplemented with 5% FBS with 42D^ENZR^ and 42F^ENZR^ cells get an additional 10µM of Enzalutamide. DU145 and VCaP were cultured in DMEM supplemented with 5% FBS *(Thermo – 26140079)*. NCI-H660 and LASPC-1 cells were grown in HITES media as per previous protocols^13,14^.

**Cell line generation and transfection:**

*42D^ENZR^* *rtA-eSpCas9*: 42D cells stably expressing tet-inducible Cas9 *(Vectorbuilder)* were created using piggybac transposon system. In short, cells were transfected with 2:1 ratio of donor plasmid (rTA-eSpCas9) to pHypbase transposase plasmid using Mirus TransIT 20/20 transfection reagent overnight. Media was changed the following morning and 5 days later, cells were subject to Puromycin *(Thermo - A1113803)* selection for 1 week.

*42D^ENZR^ BRN2^-/-^*: 42D cells expressing tet-inducible Cas9 (*42D^ENZR^ rtA-eSpCas9*) were transiently transfected with purified gRNA targeting exon 1 of BRN2 as previously used^15^. In short, cells were transfected with 2µg of purified gRNA per well of 6 well plate using Lipofectamine 3000 as per manufacturer’s protocol for RNA transfection. Next morning, media was changed to RPMI 5% FBS +10µM Enzalutamide + 1nM Doxycycline. Cells were collected for protein and mRNA 5 days after media change.

22RV1 BRN2^-/-^: Same method as 42D^ENZR^ above.

siRNA: Luciferase positive 42D^ENZR^ cells were transfected overnight with 10nM of siRNA targeting BRN2 (*Santa Cruz - sc-29838*) using Lipofectamine 3000 as per manufacturer’s protocol for siRNA transfection. Next morning, media was changed to RPMI 5% FBS +10µM Enzalutamide and 48 hours later cells were collected for Luciferase assay or Western Blot.

**Cell Proliferation:**

For adherent cell line models (including organoids), proliferation was conducted with Crystal Violet (CV). In short, 5000 for fast growing models, (16D^CRPC^, 22RV1, LNCaP, LASPC1, PC3, DU145, C4-2) or 8000 (42D^ENZR^, 42F^ENZR^) cells were counted (*BioRad Cellcounter*) and plated per well of 96 well plate. Starting cell numbers are determined with the goal of untreated wells ending with approximately 90-100% confluence at the end of the 72h experiment. Approximately 12-16h later (next morning) cells were treated with indicated doses of B18-94 for a total of 72 hours prior to fixing with 1% Glutaraldehyde and staining with CV solution (0.2% crystal violet, *Sigma - C0775*) in 10% methanol. Following staining, the CV was unstained with 10% acetic acid *(Sigma - A6283)* solution and absorbance quantified at 590nm. For non-adherant cell lines, namely LASPC1 and NCI-H660 we used CelltitreGLO *(Promega - G7570)* as per manufacturer’s protocol.

Time course experiments were conducted on Incucyte S3 Live cell imaging system. Proliferation was quantified based on confluence with the software trained to recognize each cell-line individually.

**Western Blot:**

Proteins were extracted from cells using RIPA lysis buffer with 1x PhosStop *(Roche – 4906845001)* and 1x Protease Inhibitor (*Roche - 11697498001*). Approximately 40μg of protein quantified by bicinchoninic acid (BCA assay, *Thermo – 23225*) was separated through an SDS-PAGE gel followed by wet transfer onto a PVDF membrane *(Millipore - IPVH00010)* at 80 volts for 2 hours in the cold room (4 ̊C). Membrane was blocked in 5% milk TBS-tween (0.05% tween) solution and incubated with primary antibody overnight. Next morning, after 1 hour in secondary antibody, the membranes were imaged using Licor Odyssey 9120 Imaging system with the Licor Image Studio Software. Antibody details listed below:

| **Name** | **Dilutions** | **Supplier** | **Catalogue** |
| --- | --- | --- | --- |
| BRN2 (Rb) | 1:1000 | Cell Signaling | 12137S |
| BRN2 (Ms) | 1:500 | Santa Cruz | sc-393334 |
| NCAM1 | 1:1000 | Cell Signaling | 99746S |
| PEG10 (Ms) | 1:1000 | Santa Cruz | sc-36575 |
| PEG10 (Rb) | 1:1000 | Novus | NBP2-13749 |
| ASCL1 | 1:500 | BD - Biosciences | 556604 |
| SOX2 | 1:500 | Millipore | AB5603 |
| EZH2 (Rb) | 1:2000 | Cell Signaling | 5246S |
| EZH2 (Ms) | 1:2000 | Cell Signaling | 3147S |
| CHGA | 1:1000 | Abcam | ab15160 |
| ACTIN | 1:10,000 | Sigma | A2228 |
| VINCULIN | 1:10,000 | Sigma | V9131 |
| Anti-rabbit 800 | 1:10,000 | Licor | 926-32211 |
| Anti-mouse 680 | 1:10,000 | Licor | 926-68020 |

**qRT-PCR:**

RNA was extracted from cells using RNAzol reagent (*Sigma - R4533-100ML*) according to manufacturer’s protocol. cDNA was created using SuperScript IV (*Invitrogen - 18091050*) reverse transcriptase and random hexamers *(IDT - 51-01-18-26)* from 1 μg of total mRNA. Real-time PCR amplification of cDNA was performed using Taqman FastStart Master Mix (*Roche Applied Science - 4673417001*) on ViiA7 qRT-PCR machine. Cycle-times were analyzed by ΔΔCT calculation and data is presented relative to control sample. List of Taqman probes below:

| **Taqman probe** | **Supplier** | **Catalogue** |
| --- | --- | --- |
| BRN2 | Thermo Fischer | hs00271595 |
| SOX2 | Thermo Fischer | hs01053049 |
| ASCL1 | Thermo Fischer | hs00269932 |
| EZH2 | Thermo Fischer | hs00544830 |
| PEG10 | Thermo Fischer | hs01122880 |
| CHGA | Thermo Fischer | hs00154441 |
| NCAM1 | Thermo Fischer | hs00941830 |
| GAPDH | Thermo Fischer | hs02786624 |

**Luciferase:**

42D^ENZR^ cells were stably transfected with a custom BRN2 reporter assay designed with palindromes of 3 different BRN2 response elements both monomers and homodimers separated by 2 spacer nucleotides ^16-18^. Luciferase positive 42D cells were treated with indicated doses of B18-94 overnight in 12 well plates with 3 wells per dose. Next morning, cells are collected with Passive Lysis Buffer (*Promega* - #E1941) and bioluminescence was measured using Tecan Luminometer as previously described^15^. Bioluminescence values were normalized to protein concentration to adjust for any cell number variations.

*Reporter sequence:*

“TTATGTTAATTTAAATTAACATAAATGAATTAATTCATGAGTATACTCATGAATTAATTCATGAGATGAATTTTATTCATGAGCCTGGATATCCAGGCTCATGAATAAAATTCATCTCTTATGTTAATAATTATTAACATAA”

**Drug Affinity Responsive Target Stability (DARTS):** NP40 lysis buffer and TNC buffer were prepared according to original protocol from Pai et al.^19^. Pronase was purchased from (*Sigma* 7433-2).

*From cell lysate:* Whole cell lysate from 42D^ENZR^ cells at 4-5 µg/µL concentration is incubated with B18-94 (6µM) or matched volume of DMSO for ~1 hour. Samples are equally divided into 30uL aliquots of DMSO or B18-94 with each pair receiving 2 µL of each pronase dilution from a stock of 1.25mg/mL serving as 1:100 dilution. 30 minutes after adding Pronase dilutions, the reaction is ended with addition of 10 µL of 4x SB (+βMe) and boiled at 100 ̊C for 5 minutes. Samples were then subject to WB and immunostaining with BRN2 antibody.

*From purified protein:* Purified BRN2-DBD either WT, T306A or V400E mutants were incubated with B18-94 (6µM) or matched volume of DMSO for ~1 hour. Samples are equally divided into aliquots containing 20ug of purified protein plus DMSO or B18-94 with each pair receiving 2 µL of each Pronase dilution from a stock of 1.25mg/mL serving as 1:100 dilution. 30 minutes after adding Pronase dilutions, the reaction is ended with addition of 10 µL of 4x SB (+βMe) and boiled at 100 ̊C for 5 minutes. Samples were separated through and SDS-PAGE gel and stained with Brilliant Blue protein dye.

**Microscale Thermophoresis (MST):** Purified BRN2-DBD, both WT and mutants are labeled with Monolith Protein Labeling Kit RED-NHS according to manufacturer’s protocol to a final concentration of 2µM. Labeled protein and B18-94 are diluted in PBS-Tween (0.05%) + 3% DMSO to a final concentration of 10nM and 100µM respectively. Aliquots of 10nM labeled protein was combined in 1:1 ratio with 100µM of B18-94 and serially diluted by a ratio of 1:2 in subsequent aliquots. Samples were loaded into Nanotemper Monolith Premium Capillaries and subject to MST analysis on Nanotemper Monolith NT.115 PICO Red machine.

**RNA extraction, sequencing and analysis:**

Total RNA was extracted using RNAzol (*Sigma - R4533*) using manufacture’s protocol.

*RNA-seq:* Library construction was done using NEBnext Ultra ii Standard RNA library prep kit followed by sequencing on Illumina NextSeq500. For RNA sequencing of cell lines, data was de-multiplexed using bcl2fastq2 Conversion Software (version 2.20) and the resultant read sequences were aligned to the hg19 human reference genome using STAR aligner ^20^. Assembly and differential expression were estimated using Cufflinks software (version 2.2.1) ^21^ available through the Illumina BaseSpace Sequence Hub. Gene counts (FPKM) were normalized using DESeq2 ^22^ and subsequently log-transformed.

*Microarray:* Samples preparation was done for Agilent’s One-Color Microarray-Based Gene Expression Analysis Low Input Quick Amp Labeling v6.0. 100 ng of total RNA was used to generate Cyanine-3 labeled cRNA as an input. Samples were then hybridized on Agilent SurePrint G3 Human GE 8 × 60 K Microarray (AMDID 028004) which were then scanned with the Agilent DNA Microarray Scanner at 3 µm scan resolution. Data was processed with Agilent Feature Extraction 11.0.1.1 and processed/quantile normalized with Agilent GeneSpring 12.0.

**Sub-cellular Fractionation:**

Thermo Fisher fractionation kit used according to manufacturer’s protocol (*cat. #78840*). In short, ~10 million cells treated with increasing doses of B18-94 for 16 hours were harvested and sequentially incubated through cytoplasmic extraction buffer (500µL), membrane extraction buffer (500µL) and nuclear extraction buffer (250µL) to collect the cytoplasmic, membrane and nuclear soluble fractions, respectively. Remaining pellet was incubated with CaCl_2_ and Micrococcal Nuclease for extraction of chromatin fraction (250µL). Cytoplasmic, nuclear soluble and chromatin fractions were analyzed for indicated proteins using Western Blot protocol.

**ChIP-seq and analysis:**

Approximately 15 million 42D^ENZR^ and NCI-H660 cells were fixed with 1% Formaldehyde for 10 mins at RT and quenched with 0.125M Glycine for 5 mins. Cell pellets were lysed with cell and nuclear lysis buffers provided in Millipore Magna ChIP kit *(#17-10085)*. Nuclear lysate was sonicated with Covaris M220 sonicator at high power for 6 minutes to get average size of 250-450 base pairs. Lysate spun down to remove debris. 75μL of sonicated nuclear lysate was diluted with 425μL of ChIP dilution buffer from Magna ChIP kit and incubated overnight with 20μL of magnetic A/G beads plus 5μL of BRN2-N-terminus antibody *(Cell Signaling - #12137)* and 3μg of BRN2-C-terminus antibody *(Santa Cruz - sc-393324)*. The following morning, samples were washed, treated with Proteinase K and reverse-crosslinked as per manufacturer’s instructions. DNA was purified with columns provided in the Magna ChIP kit and submitted for ChIP-seq.

Sequencing libraries were constructed using the KAPA HyperPrep Kit *(Roche #KK8504)* with Illumina TruSeq indexes (Illumina - RS-122-2001). Libraries were assessed by gel electrophoresis and purification followed by SPRI beads (*AMpure* - A63880) selection to remove primer dimers. Samples were quantified using the KAPA Library Quantification Kit for Illumina *(Roche #KK4824)* and sequenced using the Illumina NextSeq 500. FASTQ files were aligned to the hg38 reference genome using BWA-MEM software (version 0.7.17) with default parameters.

Alignments with mapping quality less than 60 were filtered out (leaving only uniquely mapped reads) using Samtools software (version 1.9). Peak calling on .bam files was done using EaSeq’s ^23^ Adaptive Local Threshold (ALT) method with p and q-value = 1E-2 and fold change of 2. Peaks were annotated to either the 5’ or 3’ end of the closest gene body. Pathway analysis on list of genes was done using gProfiler ^24^ with C2 and C5 MSigDB datasets ^25,26^.

**Gene Set Enrichment Analysis and downstream Biological Clustering:**

Gene-set enrichment analysis was conducted with GSEA desktop application from the Broad Institute. Pathways with nominal *p* value < 0.05 and false discovery rate (FDR) < 0.25 were considered significantly enriched. Data output was fed into the analysis pipeline for the EnrichmentMap ^27^ package in Cytoscape ^28^ with a similarity cut-off of 0.4 and p-value cut-off of 0.05. Each node is 1 pathway from C2 MSigDB dataset with size relative to the list of genes within the dataset and each node is colored red for NES > 2 and blue for NES < -2 (Normalized Enrichment Score).

**Annexin staining:**

42D^ENZR^, 22RV1 and NCI-H660 cells were plated in 10-cm dishes and treated with B18-94 at indicated concentration. After incubation for 48 h, cells were dissociated with Cell Stripper (*Corning* - # 25056CI) and stained with AnnexinV-FITC (*BD Biosciences* - #556547) and DAPI as live/dead stain. Stained cells were run through BD FACS CANTO II flow cytometer and analyzed using FlowJo. Data is presented as fold change of GeoMean fluorescent intensity.

**Statistics and data visualizations:**

Graphical visualization and statistics were performed in GraphPad Prism (V8). In all Bar graphs and data is combination of 3 or more biological replicates with statistical significance evaluated using a two-tailed student’s *t*-test within Graphpad. Analysis of Variance (Anova) was utilized for growth curves from Incucyte and *in vivo* experiments.

For both types of statistical analysis, *p*-value<0.05 was considered significant with asterisk indicating the following p-values: *,*p*< 0.05; **, *p* < 0.01; ***, p < 0.001; ****, p < 0.0001.

**In vivo studies:**

*Pharmacokinetics:* Five male Nu/Nu mice (*Envigo*) were given IP or PO dose (50mg/kg) of drug formulation. Blood was collected by tail vein bleed (~70 µl per time point per mouse; see SOP) in different time points (n=8) after administration: 0, 0.25h, 0.5h, 1h, 2h, 4h, 8h, 24h.Mice will be anesthetized with Isofluorine before termination via exposure to CO_2_ for 90 seconds followed by cervical dislocation. Blood samples will be centrifuged at 10000 rpm, at 4°C for 5min. The serum sample will be removed with a sterile plastic transfer pipette to the Eppendorf tubes and stored at -20°C until the LC-MS/MS analysis.

*Tumor growth:* Castrated male athymic mice (Envigo) were injected subcutaneously with 1 x 10^6^ 42D^ENZR^ or NCI-H660 cells (suspended in 0.1 mL Matrigel; *BD Biosciences*). Once tumors were 100mm^3^, mice were randomly assigned to (i) vehicle (PEG400), (ii) B18-94 (indicated doses) and treated orally 5 times per week. Tumor volume measurements were performed twice weekly with digital calipers. After sacrifice, tumors were harvested and processed as follows:

RNA – tumors frozen in RNALater followed by TriZol extraction

Protein – tumors lysed with T-PER extraction buffer with protease inhibitor cocktail.

Immunohistochemistry – Formalin fixation followed by ethanol storage. Cores are embedded in paraffin. Two sections per tumor were used to create a tissue micro-array and stained for BRN2 (1:50), Ki67 (1:100), Sox2 (1:50), PEG10 (1:50), CASP3 (1:100).

All animal procedures were performed according to the guidelines of the Canadian Council on Animal Care and appropriate institutional certification.
